# Evolutionary rescue through mutation- versus recombination-rate evolution

**DOI:** 10.64898/2026.10.04.756394

**Authors:** Alexander Longcamp, Amy L. Dapper

## Abstract

Evolutionary rescue occurs when genetic adaptation prevents extinction of a declining population. Mutation and recombination generate adaptive variation, but when their rates at adaptive loci are too low, rescue may require evolution toward higher rates of genetic change. Here, we derive and compare analytical approximations for the probability of rescue through two alternative routes: mutation-rate and recombination-rate evolution. Our model comprises two resident genotypes from which the same adaptive haplotype can arise along either route, with rescue requiring the appearance of an unlinked modifier allele that increases mutation at, or recombination between, two fitness-determining loci. We show that resident genotype composition determines the relative success of the two routes. Mutation-rate evolution yields a higher rescue probability when one resident genotype is sufficiently more common because mutation can act on either genotype whereas recombination requires both. Otherwise, the two routes yield similar rescue probabilities. Even when either route makes rescue appreciably likely, however, the modifier alleles facilitating rescue are often lost before rescue is complete. This loss is driven by multiple factors: segregation can separate the modifier from the adaptive haplotype; the fitness cost of modifier expression raises extinction risk for modifier-bearing lineages; and continued modifier-induced mutation disrupts the adaptive haplotype after it forms. The latter effect can make recombination-rate modifiers more likely to persist even when mutation-rate evolution yields a higher rescue probability. Overall, these results show that both mutation- and recombination-rate evolution can induce rescue, but successful rescue need not entail persistence of the modifier allele that facilitated it.

## 1 Introduction

When facing environmental stress, declining populations may avoid extinction through genetic adaptation. This process, known as evolutionary rescue (ER; Gomulkiewicz & Holt, 1995), can be either beneficial or detrimental to native populations depending on the context. On the one hand, ER can help such populations persist; for instance, Anstett et al. (2026) tracked populations of the scarlet monkeyflower (*Mimulus cardinalis*) in California and Oregon for over a decade and found that, while several went locally extinct due to extreme drought, others recovered through rapid adaptation via alleles favored in dry environments. On the other hand, ER can protect pathogens, pests, and invasive species, as exemplified by pathogen populations evolving antibiotic resistance (Gifford et al., 2023), and by the Australian population of the invasive European rabbit—which competes with many native Australian species (Lees & Bell, 2008)—evolving resistance to the fatal biocontrol agent, the Myxoma virus (Marshall & Fenner, 1958). Characterizing the conditions that facilitate ER can therefore help guide strategies in species conservation and the management of pests and infectious disease.

Since ER depends on the establishment of adaptive genotypes, it requires a mechanism by which adaptive genetic variation is produced. Two primary processes generate such variation: mutation introduces novel alleles into the population (Orr & Unckless, 2008), whereas recombination reshuffles these alleles to produce novel genotypic combinations (Uecker & Hermisson, 2016). Both tend to generate heritable changes during gametogenesis, with mutation occurring through errors in DNA replication and repair and recombination through crossing over. Both can also, in principle, impose fitness costs even when they tend to generate adaptive variation at focal loci: mutation can produce deleterious mutations elsewhere in the genome, whereas recombination can disrupt favorable genetic associations. A key distinction between mutation and recombination, however, is that the former can generate offspring genotypes that differ from those of the parents regardless of the parental genotypes involved, whereas recombination generates novel genotypes only when parental genotypes differ at multiple loci.

Given this dependence of ER on genetic variation, a key question is whether ER can occur through evolution toward higher mutation or recombination rates at loci contributing to adaptation. Empirical evidence suggests that both rates are heritable traits that can evolve in response to selection (Aggarwal et al., 2015; Gifford et al., 2023). For example, microbial populations evolving under drug pressure have been shown to acquire elevated mutation rates that facilitate ER in the form of multidrug resistance (Gifford et al., 2023). Similarly, laboratory populations of *Drosophila melanogaster* have been found to evolve higher recombination rates when undergoing adaptation to stress in the form of desiccation and hypoxia (Aggarwal et al., 2015).

Despite these observations, theoretical work on ER has largely proceeded under the assumption that mutation and recombination rates are fixed—rather than evolving through alleles that modify them (hereafter, modifier alleles)—with most models including only mutation (e.g., Deraje & Uecker, 2026; Gomulkiewicz & Holt, 1995; Nyhoegen et al., 2024; Orr & Unckless, 2008) but some also considering recombination (Bourne et al., 2014; Kouyos et al., 2009; Uecker & Hermisson, 2016). Notable exceptions to the fixed-rate assumption include the work of Greenspoon and Mideo (2017) and Tuffaha and Wahl (2025), who showed that mutation-rate modifier alleles can facilitate ER despite imposing fitness costs. Neither study, however, considered recombination-rate evolution (Greenspoon & Mideo, 2017; Tuffaha & Wahl, 2025). Conversely, theoretical studies have examined the evolution of mutation rates (Liberman et al., 2011) and recombination rates (Otto & Michalakis, 1998) during adaptation to changing environments, and have compared the evolution of these rates under such conditions (Carja et al., 2014), but have generally not considered adaptation that saves otherwise doomed populations from extinction and thus results in ER. Consequently, still unclear is whether—all else equal (assumed henceforth)—recombination-rate evolution can provide a viable route to ER and, if so, how effectively it facilitates ER relative to mutation-rate evolution.

Here, we address this gap by deriving and comparing analytical approximations for the probability of ER through mutation- versus recombination-rate evolution. We first introduce a three-locus population genetic model comprising two loci affecting fitness (hereafter, the major loci) and an unlinked third locus controlling either mutation at those loci or recombination between them (hereafter, the modifier locus), with mutation at the modifier locus occurring at a rate distinct from mutation at the major loci. We then adapt this model for ER via the *de novo* appearance of a modifier allele that confers a higher mutation or recombination rate among the major loci. To ensure that the population goes extinct without this focal modifier allele, we assume that the background rates of genetic change involving the major loci are negligible. Furthermore, following classical ER theory in which rescue proceeds through a single mutation (e.g., Martin et al., 2013; Orr & Unckless, 2008), the resident population is assumed to comprise two genotypes such that, once the modifier allele is present, the same adaptive genotype can be generated through a single genetic event—either a mutation or a crossover. Under these assumptions, we approximate the establishment probability of each lineage that can arise during ER and then use these per-lineage probabilities to approximate the overall probability of ER.

## 2 Model

In this section, we first introduce a three-locus population genetic model and then adapt it for ER via a modifier allele. Notation frequently used throughout the manuscript is listed in Table 1.

**Table 1:**
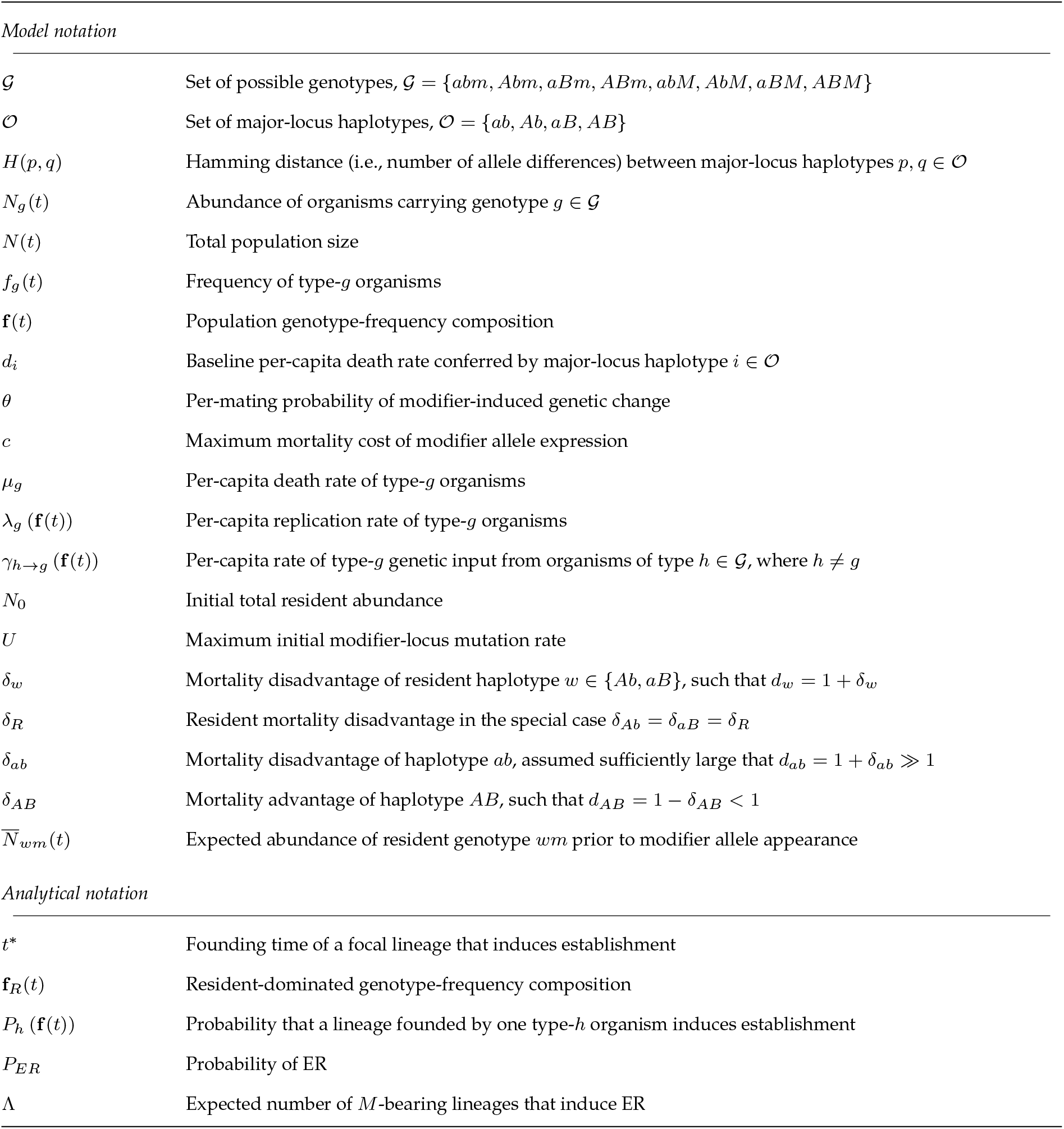
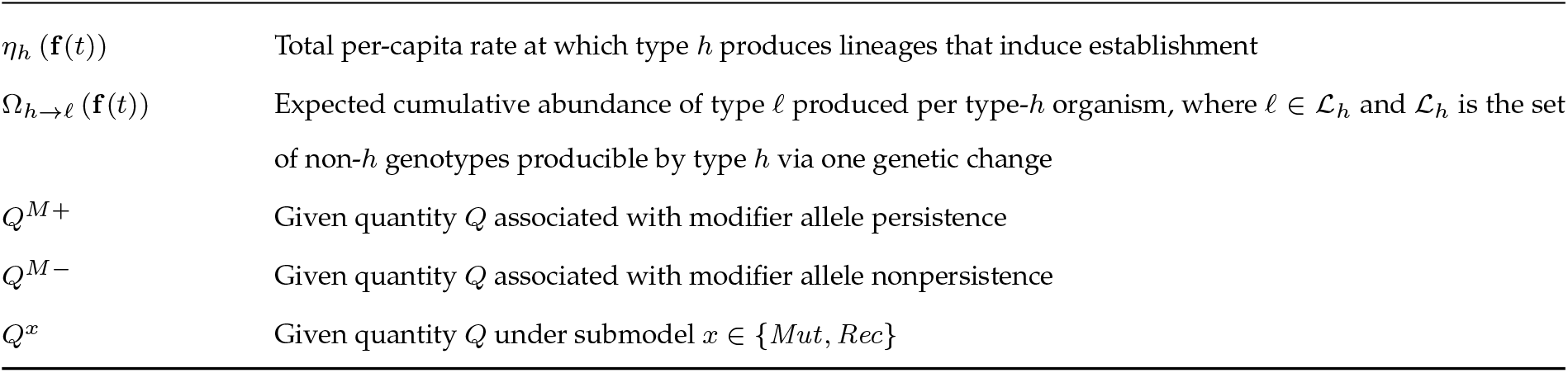
Frequently used notation.

| Symbol | Definition |
| --- | --- |
| *Model notation* |  |
| $\mathcal{G}$ | Set of possible genotypes, $\mathcal{G} = \{abm, Abm, aBm, ABm, abM, AbM, aBM, ABM\}$ |
| $\mathcal{O}$ | Set of major-locus haplotypes, $\mathcal{O} = \{ab, Ab, aB, AB\}$ |
| $H(p, q)$ | Hamming distance (i.e., number of allele differences) between major-locus haplotypes $p, q \in \mathcal{O}$ |
| $N_g(t)$ | Abundance of organisms carrying genotype $g \in \mathcal{G}$ |
| $N(t)$ | Total population size |
| $f_g(t)$ | Frequency of type-$g$ organisms |
| $\mathbf{f}(t)$ | Population genotype-frequency composition |
| $d_i$ | Baseline per-capita death rate conferred by major-locus haplotype $i \in \mathcal{O}$ |
| $\theta$ | Per-mating probability of modifier-induced genetic change |
| $c$ | Maximum mortality cost of modifier allele expression |
| $\mu_g$ | Per-capita death rate of type-$g$ organisms |
| $\lambda_g(\mathbf{f}(t))$ | Per-capita replication rate of type-$g$ organisms |
| $\gamma_{h \rightarrow g}(\mathbf{f}(t))$ | Per-capita rate of type-$g$ genetic input from organisms of type $h \in \mathcal{G}$, where $h \neq g$ |
| $N_0$ | Initial total resident abundance |
| $U$ | Maximum initial modifier-locus mutation rate |
| $\delta_w$ | Mortality disadvantage of resident haplotype $w \in \{Ab, aB\}$, such that $d_w = 1 + \delta_w$ |
| $\delta_R$ | Resident mortality disadvantage in the special case $\delta_{Ab} = \delta_{aB} = \delta_R$ |
| $\delta_{ab}$ | Mortality disadvantage of haplotype $ab$, assumed sufficiently large that $d_{ab} = 1 + \delta_{ab} \gg 1$ |
| $\delta_{AB}$ | Mortality advantage of haplotype $AB$, such that $d_{AB} = 1 - \delta_{AB} < 1$ |
| $\bar{N}_{wm}(t)$ | Expected abundance of resident genotype $wm$ prior to modifier allele appearance |
| *Analytical notation* |  |
| $t^*$ | Founding time of a focal lineage that induces establishment |
| $\mathbf{f}_R(t)$ | Resident-dominated genotype-frequency composition |
| $P_h(\mathbf{f}(t))$ | Probability that a lineage founded by one type-$h$ organism induces establishment |
| $P_{ER}$ | Probability of ER |
| $\Lambda$ | Expected number of $M$-bearing lineages that induce ER |
| $\eta_h(\mathbf{f}(t))$ | Total per-capita rate at which type $h$ produces lineages that induce establishment |
| $\Omega_{h \rightarrow \ell}(\mathbf{f}(t))$ | Expected cumulative abundance of type $\ell$ produced per type-$h$ organism, where $\ell \in \mathcal{L}_h$ and $\mathcal{L}_h$ is the set of non-$h$ genotypes producible by type $h$ via one genetic change |
| $Q^{M+}$ | Given quantity $Q$ associated with modifier allele persistence |
| $Q^{M-}$ | Given quantity $Q$ associated with modifier allele nonpersistence |
| $Q^x$ | Given quantity $Q$ under submodel $x \in \{Mut, Rec\}$ |

### 2.1 Population genetic model

We model a well-mixed, randomly mating population that changes according to a continuous-time birthdeath process, such that state variables change in increments of Δ ∈ {−1, +1} over infinitesimal time intervals (*t, t* + *dt*). The population is assumed to have a large enough carrying capacity to be negligibly affected by competition. Organisms are haploid, with reproduction occurring through the fusion of two haploid gametes into a transient diploid zygote that undergoes meiosis to produce four haploid products. Two of these products are always identical to the parental gametes and are consequently interpreted as replacing the parental gametes, while the remaining two are interpreted as new offspring. For simplicity, only one offspring is added to the focal population per reproductive event, which can be interpreted as tracking only females in a population with a 50:50 sex ratio (Hoppensteadt, 1976): one offspring is randomly assigned to the focal female population, while the other is assigned to an unmodeled male population with the same size and genetic composition as the focal population. A given organism’s genotype is characterized by three biallelic loci: two linked major loci, *A/a* and *B/b*, that affect fitness, and a third unlinked modifier locus, *M/m*, that controls either the rate of mutation at the major loci or the rate of recombination between them. Individuals can therefore carry any of eight possible genotypes, *G* = {*abm, Abm, aBm, ABm, abM, AbM, aBM, ABM* }, each consisting of one of four major-locus haplo-types, *O* = {*ab, Ab, aB, AB*}, paired with either allele *m* or *M* . The abundance of organisms carrying genotype *g* ∈ *G* is denoted *N*_*g*_(*t*) ≥ 0; total population size is 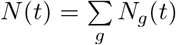; and the frequency of genotype *g* is *f*_*g*_(*t*) = *N*_*g*_(*t*)*/N* (*t*). We refer to the vector of these frequencies as the population’s genotype-frequency composition, denoted

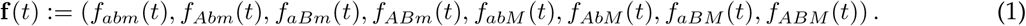

The dynamics of each genotype depend critically on the modifier locus. Whether the modifier locus controls mutation or recombination among the major loci, allele *m* induces a negligible rate of genetic change whereas allele *M* —the modifier allele—permits a non-negligible rate. Modifier-induced genetic change occurs exclusively during the brief diploid zygote phase of reproduction, prior to the production of meiotic products, and only in zygotes carrying at least one copy of allele *M*, such that allele *m* is recessive. The permating probability that an *M* -carrying zygote undergoes modifier-induced genetic change is denoted by 0 *< θ <* 1. Similar to Tuffaha and Wahl’s (2025) model of mutation-rate evolution, wherein the fitness cost of modifier allele expression is proportional to the modifier-induced mutation rate, we assume *M* carriers incur a direct adult mortality cost proportional to *θ*. Consequently, letting *T* (*N*_*g*_(*t*) → *N*_*g*_(*t*) + Δ) represent the rate of transition from state *N*_*g*_(*t*) to *N*_*g*_(*t*) + Δ during (*t, t* + *dt*), the rate of type-*g* mortality is

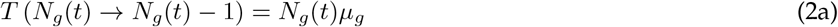

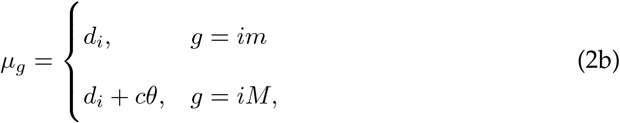

where *µ*_*g*_ *>* 0 is the per-capita death rate of genotype *g*; *d*_*i*_ *>* 0 is the baseline per-capita death rate conferred by major-locus haplotype *i* ∈ *O*; and *c >* 0 is the maximum mortality cost of *M* expression. While the production of type-*g* organisms is not directly reduced by *M* expression, it remains strongly influenced by the state of the modifier locus. One potential source of production is mutation at the modifier locus itself, whereby type-*im* organisms mutate to type-*iM* at a per-capita rate *u* ≥ 0. For simplicity, we follow previous theory (Longcamp & Draghi, 2025; Wilson et al., 2017) by assuming these mutations from *m* to *M* occur outside of reproduction and therefore independently of modifier-induced genetic change at the major loci, while remaining sufficiently rare that their effect on the abundance of the mutating genotype is negligible (i.e., a mutation from *m* to *M* does not reduce *N*_*im*_(*t*)). Back mutations from *M* to *m* are ignored. The remaining contributions to type-*g* production arise through (i) replication by type-*g* organisms and (ii) both the modifier-induced genetic change and the modifier-locus segregation that occur during reproduction between non-type-*g* organisms (hereafter, type-*g*’s genetic input). The rate of type-*g* production is thus

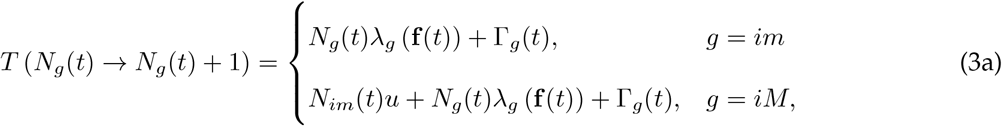

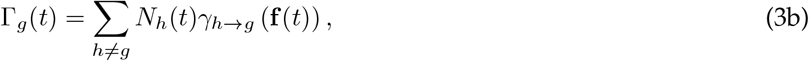

where 0 ≤ *λ*_*g*_ (**f** (*t*)) ≤ 1 is the per-capita replication rate of type-*g* organisms; Γ_*g*_(*t*) ≥ 0 is the total rate of type-*g* genetic input; and *γ*_*h*→*g*_ (**f** (*t*)) ≥ 0 is the per-capita rate of type-*g* genetic input from organisms of type *h* ≠*g*, where *h* ∈ *G*. Because mating is random, each term in *λ*_*g*_ (**f** (*t*)) is the product of the frequency of a mating partner carrying major-locus haplotype *j* ∈ *O* and the probability that a mating between carriers of haplotypes *i* and *j* produces a female type-*g* offspring. When *i* ≠*j*, the reciprocal *i* × *j* and *j* × *i* mating directions each produce a female type-*g* offspring with probability 1*/*2, giving a combined probability of one. For matings between type-*im* and type-*jm* parents, the replication probability is therefore one, since no modifier-induced genetic change occurs. When at least one parent carries allele *M*, however, replication can be disrupted by modifier-induced mutation or recombination and, when the parents differ at the modifier locus, by modifier-locus segregation. The corresponding replication probability is denoted by 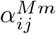 when the parents differ at the modifier locus and by 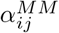 when both carry *M*, yielding

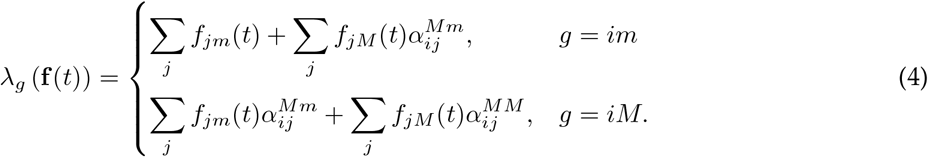

The genetic-input rate Γ_*g*_(*t*) is derived similarly, except that each contribution is the product of a non-*g* parental abundance, a non-*g* mating-partner frequency, and the corresponding probability of producing type *g*. One source of input occurs when one parent carries haplotype *i* but the opposite modifier-locus allele from type *g*, while the other carries the focal modifier-locus allele but a nonfocal major-locus haplotype *j* ≠*i*. Because these parents differ at the modifier locus, this contribution is governed by the same 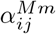 used in *λ*_*g*_ (**f** (*t*)). Alternatively, when both parents carry nonfocal major-locus haplotypes *j, k* ≠*i*, producing *g* requires modifier-induced mutation or recombination to generate haplotype *i*. We define 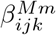 and 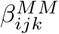 as the per-mating-direction probabilities of type-*g* production when the parents carry *M* and *m*, or when both parents carry *M*, respectively. Therefore,

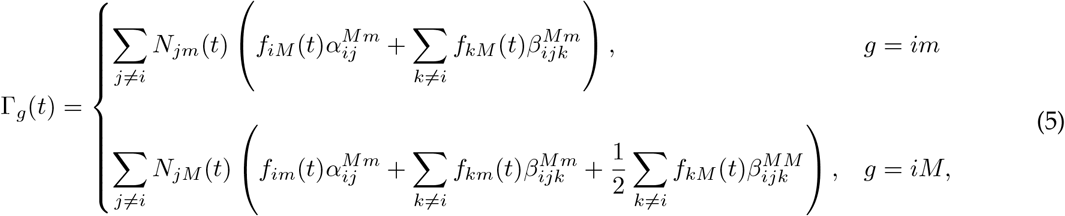

where the factor 1*/*2 in the *M* × *M* term gives the probability that the generated type-*g* organism is the offspring assigned to the focal female population. For *M* ×*m* matings, the two reciprocal mating directions are combined within each term, so their factor of two cancels this probability of 1*/*2. The specific expressions for coefficients 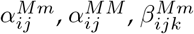, and 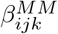 depend on whether allele *M* modifies mutation or recombination among the major loci. Accordingly, two submodels are considered: a mutation-rate modifier submodel and a recombination-rate modifier submodel. Within each submodel, the coefficients are written in terms of the Hamming distance (i.e., the number of allele differences) between pairs of major-locus haplotypes *p, q* ∈ *O*, denoted by *H*(*p, q*).

#### 2.1.1 Mutation-rate modifier

For the mutation-rate modifier submodel, the major loci are assumed to be fully linked such that the rate of recombination between them is zero. During a mating involving at least one *M* -carrying parent, however, the zygote can experience a single mutation at one of the two major loci with probability *θ >* 0, with forward and back mutations equally possible. When a mutation occurs, both the parental gamete receiving the mutation and the mutated locus are chosen at random. In Appendix A, we derive the coefficients that arise under these assumptions; the probabilities of type-*g* replication via *M* × *m* and *M* × *M* matings are

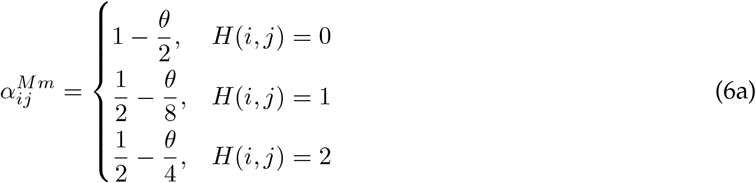

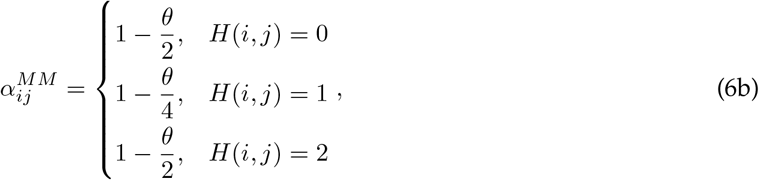

and the probability that mutation generates type *g* from nonfocal parental haplotypes *j, k* ≠*i* in a given *M* × *m* or *M* × *M* mating direction is

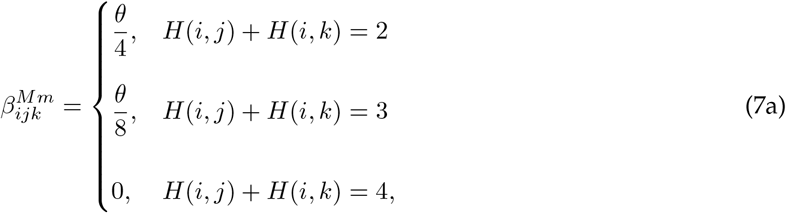

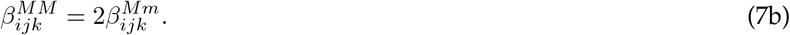

Equation (6) shows that type-*g* replication depends on the Hamming distance between the parental haplotypes, *H*(*i, j*). In *M* × *m* matings, 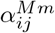 has a baseline value of one when the parental haplotypes are identical, whereas it has a baseline value of 1*/*2 when they differ because modifier-locus segregation causes the focal modifier allele to be inherited with haplotype *i* only half as often. Mutation generally reduces these probabilities by disrupting haplotype *i*, although this effect can be partly offset when the alternative parental haplotype is one mutation away from *i* and can therefore regenerate it. The same general dependence applies to 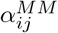, except that modifier-locus segregation no longer limits replication because both parents carry allele *M* . Similarly, Eq. (7) shows that genetic input into type *g* declines as the parental haplotypes become more distant from *i* and is absent when neither can generate *i* by a single mutation. Because *M* × *M* matings do not require haplotype *i* to segregate with allele 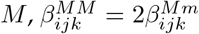.

#### 2.1.2 Recombination-rate modifier

For the recombination-rate modifier submodel, we assume the rate of mutation at the major loci is zero. In contrast, a zygote carrying allele *M* experiences a crossover between the major loci with probability *θ >* 0. We derive the coefficients comprising this submodel in Appendix B; the probabilities of type-*g* replication via *M* × *m* and *M* × *M* matings are

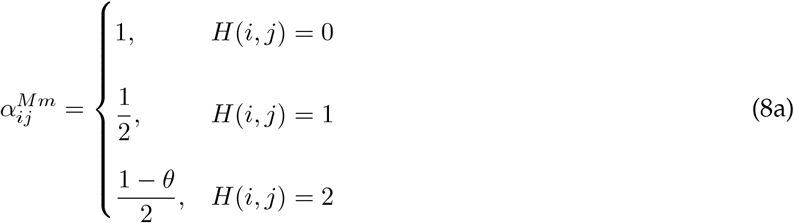

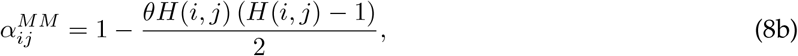

and the per-mating-direction probabilities of recombination generating type *g* from nonfocal parental haplotypes *j, k* ≠*i* are

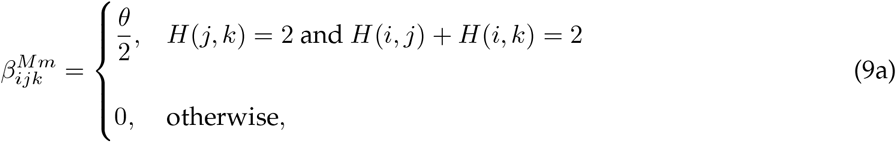

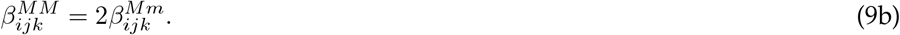

Equation (8) shows that recombination differs from mutation in having more restricted effects on type-*g* replication. As under mutation-rate modification, 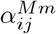 equals one when the parental haplotypes are identical and otherwise has a baseline value of 1*/*2 owing to modifier-locus segregation. Recombination alters this baseline only when the parental haplotypes differ at both major loci, *H*(*i, j*) = 2, because only then can a crossover disrupt haplotype *i*. The same restriction applies to 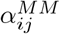, with the modifier-locus segregation constraint again absent because both parents carry allele *M* . Equation (9) shows that recombinationmediated genetic input is similarly restricted: type *g* can be generated only when the parental haplotypes differ at both major loci and *i* is one of the resulting recombinant haplotypes. As in the mutation-rate modifier submodel, 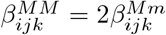 because *M* × *M* matings do not require haplotype *i* to segregate with allele *M* .

### 2.2 ER via a modifier allele

We now adapt our population genetic model for ER via the modifier allele *M* . At the onset of environmental change, *t* = 0, the population contains only type-*Abm* and type-*aBm* organisms (hereafter, residents), with a large total abundance, *N*_0_, such that *N*_*Abm*_(0)+*N*_*aBm*_(0) = *N*_0_ ≫ 1. For simplicity, the initial abundance of each resident genotype is treated as a historically determined condition arising from processes not explicitly modeled. Following the environmental change, each major-locus haplotype *i* experiences a change in its baseline per-capita death rate of magnitude *δ*_*i*_ *>* 0, with mortality disadvantages increasing the baseline death rate by *δ*_*i*_ and mortality advantages decreasing it by *δ*_*i*_. For each resident haplotype *w* ∈ {*Ab, aB*}, we assume a mortality disadvantage, such that *d*_*w*_ = 1 + *δ*_*w*_ *>* 1, with *δ*_*Ab*_ = *δ*_*aB*_ = *δ*_*R*_ in the special case of equal resident disadvantages. Consequently, since type-*wm* residents have a per-capita replication rate of *λ*_*wm*_(**f** (*t*)) = 1 when organisms carrying *M* are absent, their expected abundance under such conditions is

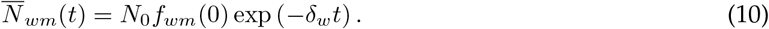

During this decline, *M* can arise through *de novo* mutation, forming an *AbM* or *aBM* mutant. We let the per-capita modifier-locus mutation rate be *u* = *U/N*_0_, so that the total rate of modifier-locus mutation, *N* (*t*)*u*, has a small maximum initial value of 0 ≤ *U <* 1, ensuring that ER does not become effectively certain due to *N*_0_ ≫ 1. Following the appearance of *M*, carriers of the *ab* and *AB* haplotypes can arise through mutation or recombination. We assume that haplotype *ab* confers a strong mortality disadvantage, such that *d*_*ab*_ = 1 + *δ*_*ab*_ ≫ 1 and thus *ab* carriers have strongly negative expected growth rates. Haplotype *AB*, in contrast, confers a mortality advantage, such that *d*_*AB*_ = 1 − *δ*_*AB*_ *<* 1 and so—while *M* remains sufficiently rare—type-*ABm* organisms have positive expected growth rates. Type-*ABM* organisms likewise have positive expected growth rates under such conditions provided that *δ*_*AB*_ is adequately large and the cost of *M* expression, *c*, is sufficiently low. Thus, ER requires the appearance and subsequent establishment of the *AB*-carrier population, with establishment occurring when *AB* carriers reach an abundance of *N*_*ABm*_(*t*) + *N*_*ABM*_ (*t*) ≥ *N*_0_. For simplicity, the same establishment criterion is applied specifically to type-*ABM* organisms, such that their population is considered to have established, and *M* is therefore considered to have persisted, only if it reaches an abundance of *N*_*ABM*_ (*t*) ≥ *N*_0_.

## 3 Methods

We analytically approximate the probability of ER, *P*_*ER*_, in each submodel and use stochastic simulations to test the resulting predictions. The first part of this section presents the general theory used to derive these approximations, and the second describes the simulation algorithm.

### 3.1 Theory

We classify *AB* establishment (hereafter, establishment) according to whether modifier allele *M* persists, denoted *M* +, or does not persist, denoted *M* −, with *Q*^*M*+^ and *Q*^*M*−^ denoting the corresponding values of a given quantity *Q*. The overall probability of ER can therefore be decomposed as

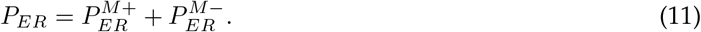

To approximate Eq. (11), we assume that mutations from *m* to *M* are sufficiently rare for the resulting *M* - bearing mutants to arise, reproduce, and die independently of one another. The number of *M* mutations that successfully induce ER prior to resident extinction is then Poisson-distributed, with an expectation denoted by Λ = Λ^*M*+^ + Λ^*M*−^. Accordingly, *P*_*ER*_ is the complement of the probability that no *M* -bearing lineage arises and succeeds, (Λ)^0^ exp (−Λ) */*(0!) = exp (−Λ), yielding

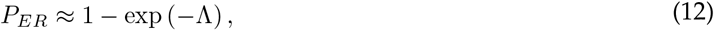

and

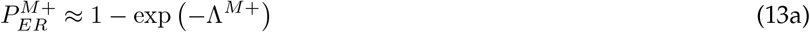

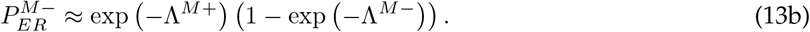

For the approximation of Λ, we use the expected resident abundances in Eq. (10), 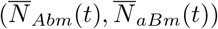, to approximate mutational input during the resident decline. Because mutants are assumed sufficiently rare to arise independently, and an *M* mutation can arise at any time *t* = *t*_*M*_ *>* 0, this gives

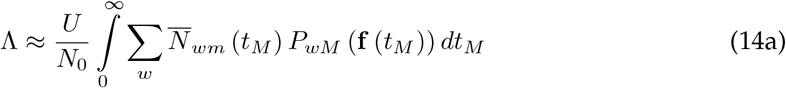

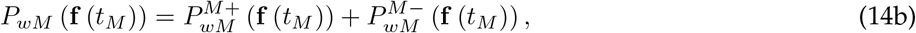

where 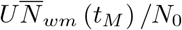 is the expected rate at which residents carrying haplotype *w* ∈ {*Ab, aB*} mutate to *wM* at time *t*_*M*_, and *P*_*wM*_ (**f** (*t*_*M*_ )) is the probability that the resulting mutant lineage ultimately produces an established *AB* population given genotype-frequency composition **f** (*t*_*M*_ ). The latter probability depends on the probability that a lineage founded at some time *t*^∗^ ≥ *t*_*M*_ by one type-*ABm* organism induces establishment, *P*_*ABm*_ (**f** (*t*^∗^)), and the analogous probability for a lineage founded by one type-*ABM* organism, *P*_*ABM*_ (**f** (*t*^∗^)), which can be written as

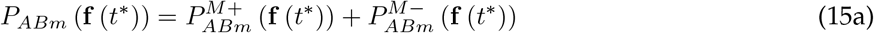

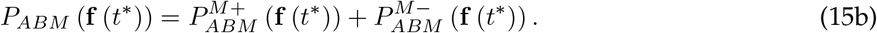

The probabilities in Eqs. (15) and (14b) can be treated as special cases of a general probability that a type-*h* lineage founded at time *t*^∗^ induces establishment, *P*_*h*_ (**f** (*t*^∗^)), where *h* ∈ {*AbM, aBM, ABm, ABM* }. To approximate this probability, we assume that (i) the focal type-*h* organisms are sufficiently rare that their own frequency and their rate of mutation at the modifier locus are negligible prior to establishment; and (ii) over a given time interval, all nonfocal genotype frequencies remain effectively constant. In Appendix C, we show that these assumptions permit a branching-process approximation:

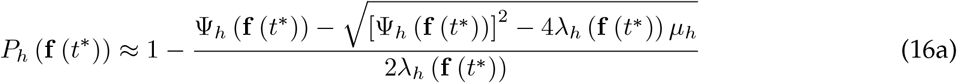

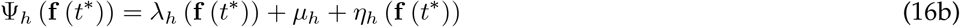

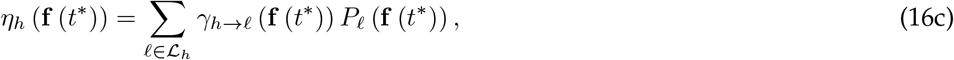

with *λ*_*h*_ (**f** (*t*^∗^)) being the per-capita type-*h* replication rate; *µ*_*h*_ the per-capita type-*h* death rate; *L*_*h*_ the set of non-*h* genotypes that can be produced by type *h* via one genetic change; *γ*_*h*→ℓ_ (**f** (*t*^∗^)) the per-capita rate at which type *h* gives rise to type ℓ ∈ *L*_*h*_; and *η*_*h*_ (**f** (*t*^∗^)) the total per-capita rate at which type *h* produces rare lineages that successfully induce establishment. The quantity Ψ_*h*_ (**f** (*t*^∗^)) is therefore the total rate of events affecting whether a rare type-*h* lineage induces establishment. Because type *h* can contribute to establishment through transient type-ℓ lineages that it produces, we additionally consider the expected abundance accumulated by such lineages before extinction. If type ℓ is deleterious, such that *µ*_ℓ_ *> λ*_ℓ_ (**f** (*t*^∗^)), this cumulative abundance is 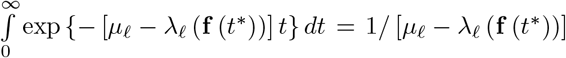. Multiplying this quantity by the rate at which type *h* produces type ℓ gives the expected type-ℓ abundance accumulated per type-*h* organism:

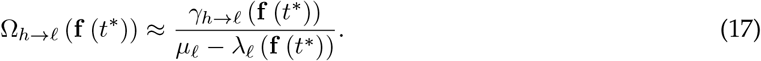

#### 3.1.1 Outline of analysis

Our forthcoming analysis proceeds stepwise toward a closed-form approximation of *P*_*ER*_. At each step preceding the final *P*_*ER*_ approximation, we apply Eq. (16) to approximate the probability that a rare focal lineage, founded at time *t*^∗^ in a resident-dominated population with genotype-frequency composition **f**_*R*_ (*t*^∗^), induces establishment. Because the focal type *h* in each *P*_*h*_ (**f**_*R*_ (*t*^∗^)) approximation is assumed to be rare, we approximate this composition as

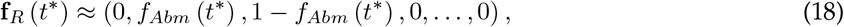

where *f*_*Abm*_ (*t*^∗^) is approximated with its expected value in Eq. (10). Given this starting composition, we first approximate the per-lineage probabilities of *ABm*- and *ABM* -induced establishment, *P*_*ABm*_ (**f**_*R*_ (*t*^∗^)) and *P*_*ABM*_ (**f**_*R*_ (*t*^∗^)), respectively. We then use the per-lineage *ABm*- and *ABM* -induced establishment probabilities to approximate each per-lineage probability of *wM* -induced establishment, *P*_*wM*_ (**f**_*R*_ (*t*^∗^)). Finally, those *P*_*wM*_ (**f**_*R*_ (*t*^∗^)) probabilities are used to approximate *P*_*ER*_. The per-capita mortality and production rates used to derive each approximation are listed in Table D1 of Appendix D.

### 3.2 Simulations

We test our analytical predictions using a tau-leaping variant of the Gillespie algorithm (Cao et al., 2006; Gillespie, 1976). To test each predicted per-lineage probability of *h*-induced establishment, *P*_*h*_ (**f**_*R*_ (*t*^∗^)), we prevent mutations from *m* to *M* by setting the maximum initial modifier-locus mutation rate to *U* = 0 and initialize residents at their expected abundances at time *t*^∗^, such that 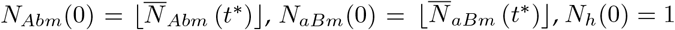 for the focal genotype *g* = *h*, and *N*_*g*_(0) = 0 for all *g* ∉ {*Abm, aBm, h*}. By contrast, to test the ER probability, we let *U >* 0 and initialize simulations with *N*_*Abm*_(0) = *N*_0_*f*_*Abm*_(0), *N*_*aBm*_(0) = *N*_0_(1 − *f*_*Abm*_(0)), and *N*_*g*_(0) = 0 for all *g* ∉ {*Abm, aBm*}. Each variable, *N*_*g*_(*t*), then changes according to its corresponding events (*N*_*g*_(*t*) → *N*_*g*_(*t*) + Δ), state changes (Δ), and transition rates (*T* (*N*_*g*_(*t*) → *N*_*g*_(*t*) + Δ)) in Eqs. (2) and (3). The size of each tau-leap, and the number of times each event occurs during the leap, are computed according to the procedure outlined in Sections IIC (excluding Step 3) and IV of Cao et al. (2006). This procedure ensures that, if *N*_*g*_(*t*) *< n*_*c*_ = 10, then all transition rates corresponding to a decrease in *N*_*g*_(*t*) are updated according to a nearly exact version of the Gillespie algorithm (Gillespie, 1976). The procedure also helps prevent *N*_*g*_(*t*) from becoming negative during a leap by approximately bounding the per-leap change in each of *N*_*g*_(*t*)’s transition rates, relative to that transition rate’s current value, by a value of *ϵ* = 0.06 (Cao et al., 2006). If *N*_*g*_(*t*) becomes negative despite this precaution, we set *N*_*g*_(*t*) = 0. We avoid an undefined leap size by adding 10^−9^ to the output of Cao et al.’s (2006) Eq. (32a) if the output equals zero, and the same modification is made to their Eq. (32b) if its output equals zero. A simulation proceeds until either all genotypes are extinct, 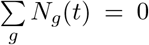, or both ER has occurred at some point— with the *AB* population having reached *N*_*ABm*_(*t*) + *N*_*ABM*_ (*t*) ≥ *N*_0_—and the fate of allele *M* has been resolved, with either all *M* -carrying organisms extinct, 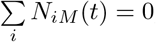, or type-*ABM* organisms established, *N*_*ABM*_ (*t*) ≥ *N*_0_. All simulations are performed using Mathematica 14.2.0.0.

## 4 Results

In this section, we derive and analyze the probability of ER, *P*_*ER*_, following the stepwise analytical approach outlined in the Methods. For the per-lineage establishment probabilities considered along the way, *t*^∗^ denotes the appearance time of the focal lineage and **f**_*R*_ (*t*^∗^) the resident-dominated genotype-frequency composition at that time. The corresponding approximations depend on the focal type’s per-capita death rate, *µ*_*h*_, replication rate, *λ*_*h*_ (**f**_*R*_ (*t*^∗^)), and rate of genetic input into type ℓ ≠*h, γ*_*h*→ℓ_ (**f**_*R*_ (*t*^∗^)). Unless stated otherwise, the two resident haplotypes, indexed by *w* ∈ {*Ab, aB*}, begin at equal frequencies, *f*_*Abm*_(0) = *f*_*aBm*_(0), and have equal mortality disadvantages, *δ*_*Ab*_ = *δ*_*aB*_ = *δ*_*R*_. For notational simplicity, we henceforth omit **f**_*R*_ (*t*^∗^) as an explicit argument unless it is needed for clarity, and let *x* ∈ {*Mut, Rec*} index the modifier submodel, with *x* = *Mut* denoting mutation-rate modification and *x* = *Rec* denoting recombination-rate modification.

### 4.1 The per-lineage probability of *ABm*-induced establishment

To begin our analysis, we consider the per-lineage probability of *ABm*-induced establishment, *P*_*ABm*_. Given our assumption that a focal lineage has a negligible modifier-locus mutation rate, the per-capita rate at which *ABm* produces lineages that later induce establishment is *η*_*ABm*_ ≈ 0. Consequently, substituting *µ*_*ABm*_, *λ*_*ABm*_, and this *η*_*ABm*_ expression into Eq. (16) gives

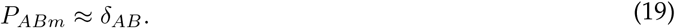

Equation (19) shows that, at least approximately speaking (assumed hereafter), the chance of an *ABm* lineage establishing increases linearly with the haplotype-*AB* mortality advantage, *δ*_*AB*_ (Figure 1). Because type *ABm* does not carry the modifier allele *M* and, while rare, effectively mates only with residents in the resident-dominated population, its per-capita replication rate remains one under both mutation- and recombination-rate modification, while its per-capita death rate decreases linearly with *δ*_*AB*_. Consequently, raising *δ*_*AB*_ reduces the chance of stochastic lineage extinction identically irrespective of modifier type.

**Figure 1:**
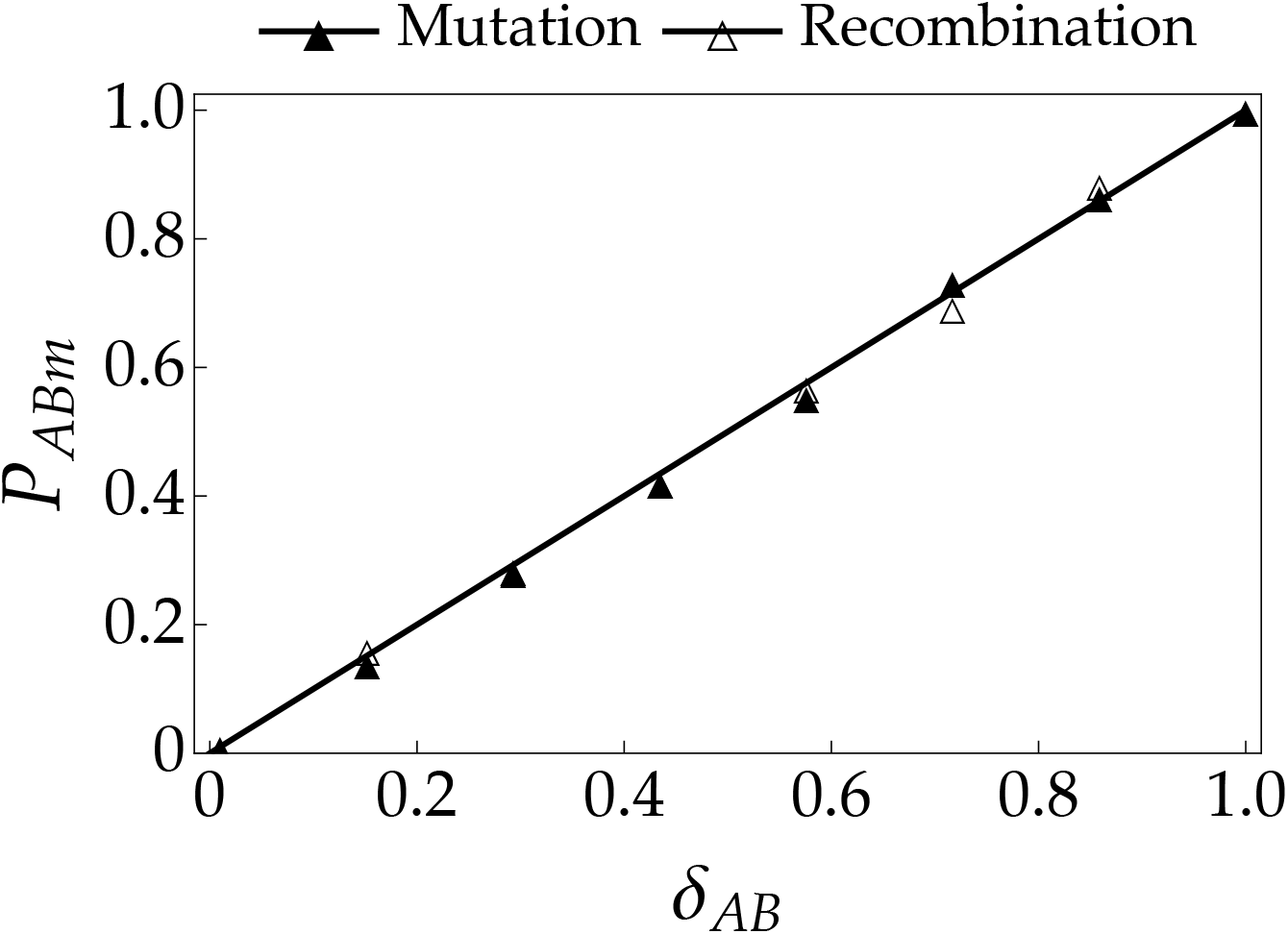
The probability of *ABm*-induced establishment, *P*_*ABm*_, as a function of the haplotype-*AB* mortality advantage, *δ*_*AB*_, under mutation-rate versus recombination-rate modification. In both submodels, *P*_*ABm*_ increases identically with *δ*_*AB*_ because type *ABm* lacks modifier allele *M* and therefore has the same replication and death rates. Markers: proportion of 3 *×* 10^3^ replicate simulations that ended in establishment; solid curve: Eq. (19). Parameter values are *c* = 0.3, *δ*_*R*_ = 0.003, *δ*_*ab*_ = 10, *N*_0_ = 10^6^, *θ* = 0.5, *t*^∗^ = 1, and *U* = 0.

### 4.2 The per-lineage probability of *ABM* -induced establishment

We next consider the per-lineage probability of *ABM* -induced establishment, *P*_*ABM*_ . We approximate this probability by accounting for establishment of the *ABM* lineage itself together with paths to establishment in which type-*ABM* organisms produce an *ABm* lineage that establishes, either directly or through a type-*wM* intermediate lineage. For simplicity, we ignore longer paths involving additional intermediate-genotype transitions, as well as paths in which type-*wM* intermediates regenerate *ABM*, which return to the focal genotype rather than provide a distinct path to *ABm* establishment. To obtain the per-capita rate of successful *ABm* production, 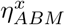, we first approximate the expected cumulative abundance of type-*wM* intermediates generated per type-*ABM* organism, 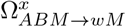. Substituting 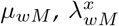, and 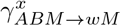 into Eq. (17) gives

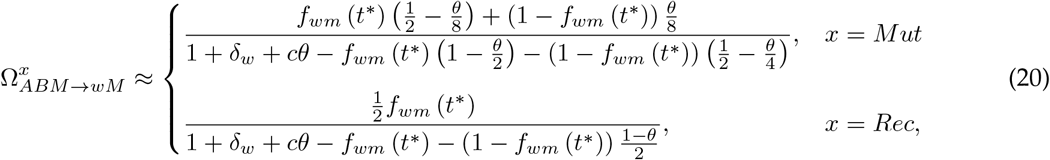

which helps determine the rate of *ABm* production per type-*ABM* organism, 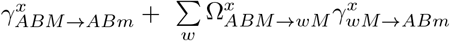. Multiplying this rate by *P*_*ABm*_ ≈ *δ*_*AB*_ (Eq. (19)) then gives the per-capita rate at which type-*ABM* organisms produce a successfully established *ABm* lineage:

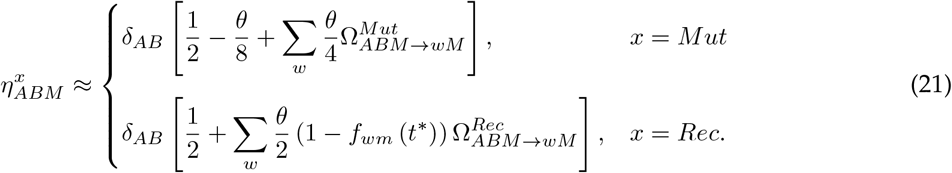

Substituting this 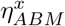 expression into Eq. (16) alongside *µ*_*ABM*_ and 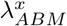, in turn yields

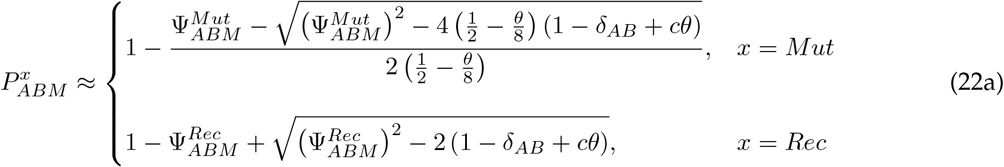

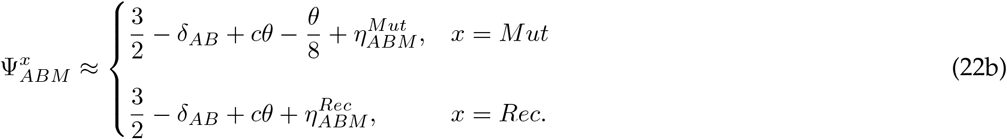

Equations (20)–(22) show that 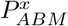 generally increases with parameter changes that reduce the mortality rates of carriers of modifier allele *M* and their descendants (Figure 2). Boosting *δ*_*AB*_ raises 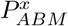 (Figure 2A) by reducing the mortality rate of type-*ABM* organisms and, through the factor *δ*_*AB*_ in Eq. (21), increasing the contribution of type-*ABm* descendant lineages that establish with probability *P*_*ABm*_ ≈ *δ*_*AB*_. Decreasing *δ*_*R*_ lowers the mortality rates of type-*wM* intermediates and increases the expected cumulative abundance of type-*wM* intermediates, 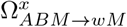, as seen through *δ*_*w*_ in Eq. (20), but causes only a comparatively weak-to-negligible increase in 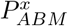 (Figure 2B) because resident mortality disadvantages act only through this intermediate-mediated path. Reducing the maximum mortality cost of modifier allele *M* expression, *c*, likewise raises 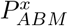 (Figure 2C) by diminishing the realized cost of *M* expression, *cθ*, which lowers the mortality rate of type-*ABM* organisms through Eq. (22) and also increases each 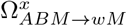.

**Figure 2:**
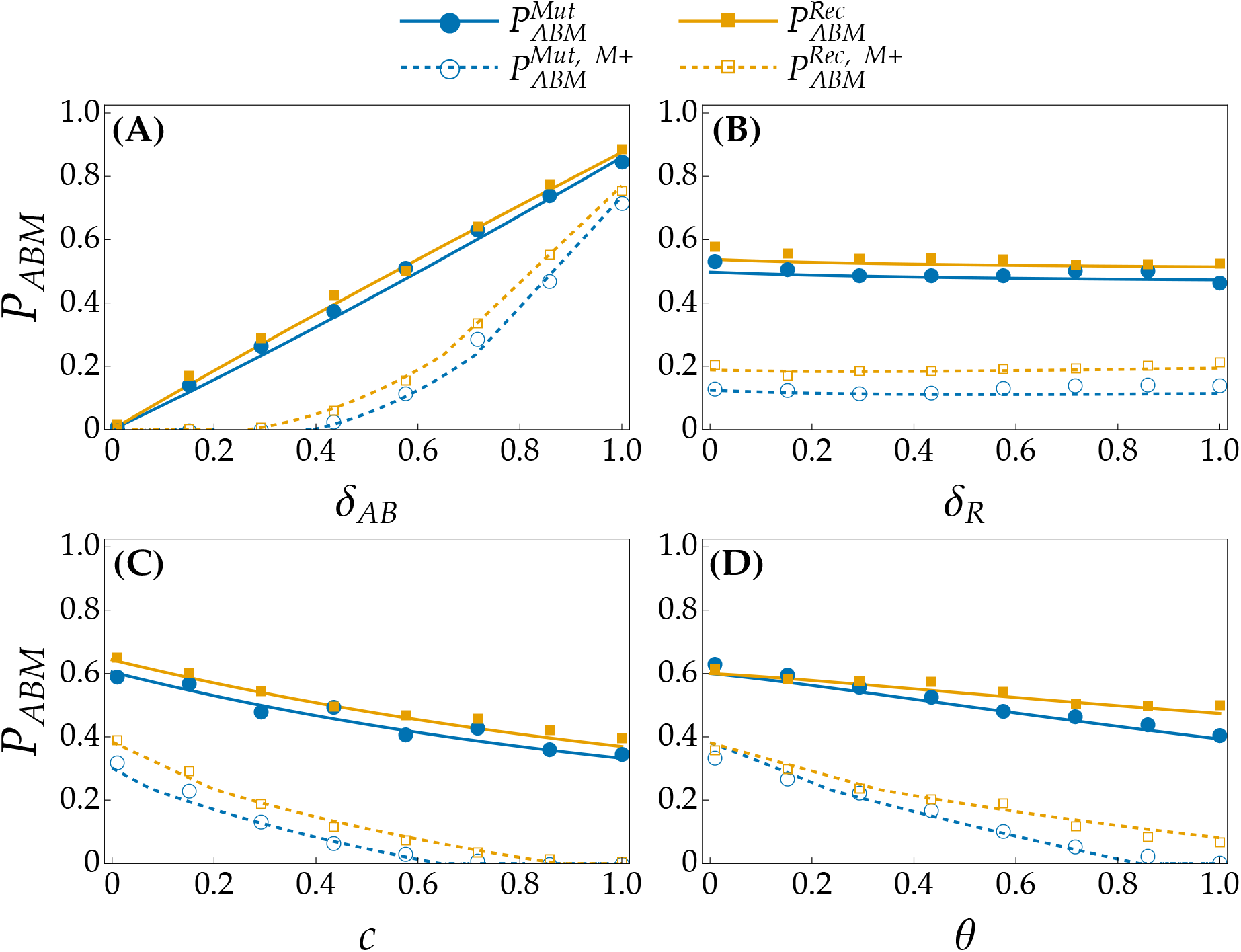
The probability of *ABM* -induced establishment, *P*_*ABM*_, and its constituent probability of *ABM* -induced establishment with modifier allele *M* persistence, 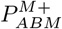, as functions of (A) the haplotype-*AB* mortality advantage, *δ*_*AB*_ ; (B) the resident mortality disadvantage, *δ*_*R*_; (C) the maximum mortality cost of *M* expression, *c*; and (D) the per-mating probability of modifier-induced genetic change, *θ*. Superscripts *Mut* and *Rec* denote the mutation- and recombination-rate modifier submodels, respectively. Each parameter affects the mortality rates of *M* -carrying organisms or their descendants, while *θ* additionally controls the rate of modifier-induced genetic change. Increasing *δ*_*AB*_ or reducing *c* raises *P*_*ABM*_ by lowering these mortality rates, whereas variation in *δ*_*R*_ has comparatively little effect because resident mortality disadvantages act only through type-*wM* intermediate lineages. Reducing *θ* likewise tends to raise *P*_*ABM*_ by lowering the mortality cost of *M* expression and, under mutation-rate modification, reducing the chance that mutation breaks apart the adaptive *AB* haplotype. As shown in each panel, recombination-rate modification can consequently yield larger *P*_*ABM*_ by better preserving *AB*. The behavior of 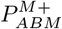 broadly resembles that of *P*_*ABM*_, but 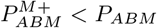 because establishment can proceed through type-*ABm* descendants after *M* is lost. Markers: proportion of 3 *×* 10^3^ replicate simulations ending in establishment; solid curves: Eq. (22); dashed curves: Eq. (27). Unless stated otherwise, *c* = 0.3, *δ*_*AB*_ = 0.6, *δ*_*R*_ = 0.003, *δ*_*ab*_ = 10, *N*_0_ = 10^4^, *θ* = 0.5, *t*^∗^ = 1, and *U* = 0.

The probability of modifier-induced genetic change, *θ*, plays a more complex role because it affects both modifier-associated mortality and genetic change at the major loci, with smaller *θ* generally raising 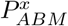 (Figure 2D) by reducing *cθ*. Under mutation-rate modification, Eq. (21) further shows that diminishing *θ* increases the per-capita rate at which *ABM* produces 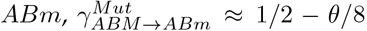, while Eq. (22) shows the corresponding increase in the per-capita type-*ABM* replication rate, 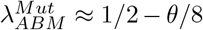, because mutation can break apart the adaptive *AB* haplotype. Under recombination-rate modification, these rates instead remain approximately 1*/*2: an *ABM* × *Abm* mating is homozygous at the *A* locus, whereas an *ABM* × *aBm* mating is homozygous at the *B* locus, so recombination between the major loci cannot break apart *AB*. Consequently, recombination-rate modification can yield an appreciably larger probability of *ABM* -induced establishment, 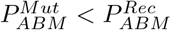 (Figure 2).

#### 4.2.1 The per-lineage probability of *ABM* -induced establishment with modifier allele persistence

Now that we have approximated 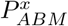, we can approximate the probability of *ABM* -induced establishment with *M* persistence, 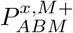. To do so, we account for the possibility that an *ABM* lineage either establishes directly or first produces an *ABm* lineage and subsequently establishes while that *ABm* lineage is transitioning from rarity toward commonality. In the latter case, we additionally consider the possibility that a type-*wM* intermediate subsequently produces a new *ABM* lineage that ultimately establishes while the *ABm* lineage is becoming common. We approximate the changing genotype-frequency composition encountered by the focal *ABM* lineage and any type-*wM* descendants it produces during this transition with an effective composition, 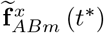, that captures both the increasing frequency of type *ABm* and the corresponding decrease in the frequencies of the resident genotypes. Specifically, we write

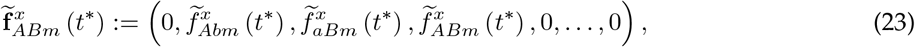

where 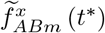 is the effective frequency of type *ABm* during the transition 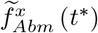 and 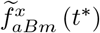 are the corresponding effective frequencies of the two resident genotypes. We derive these effective frequencies in Appendix E. Under this composition, we first approximate the expected cumulative abundance of type-*wM* intermediates generated per type-*ABM* organism, 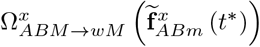. Substituting 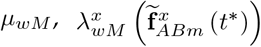, and 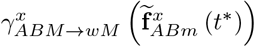 into Eq. (17) gives

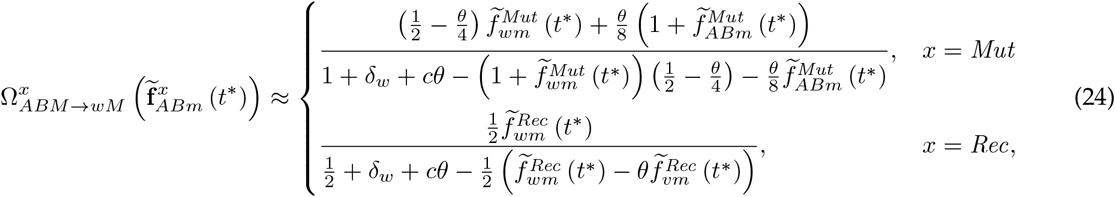

with *v* ∈ {*Ab, aB*} being the alternative haplotype to *w*, i.e., *v* ≠*w*. The per-capita rate at which type-*ABM* organisms produce a new *ABM* lineage through type-*wM* intermediates is then approximately 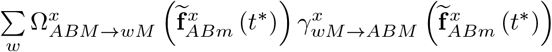. Multiplying this rate by 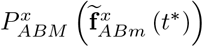, gives the per-capita rate at which type-*ABM* organisms produce lineages that successfully induce *ABM* establishment:

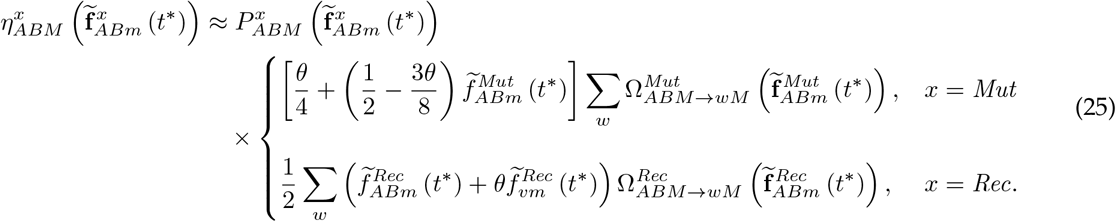

By substituting 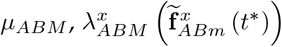, and this 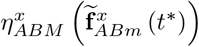 expression into Eq. (16); solving for 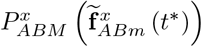; and retaining the largest solution in [0, 1], we obtain

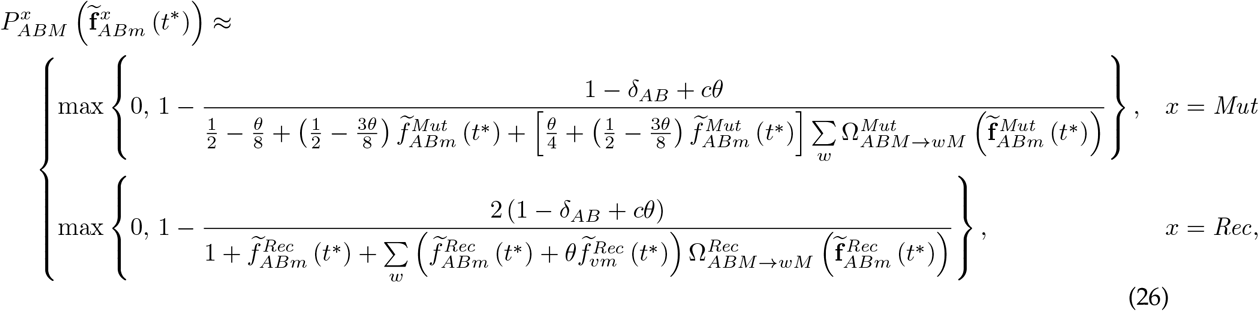

which approximates the probability that an *ABM* lineage establishes while an *ABm* lineage is growing from rarity toward commonality. Given this approximation and that of 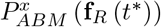 in Eq. (22), we finally approximate the probability of *ABM* -induced establishment with *M* persistence as

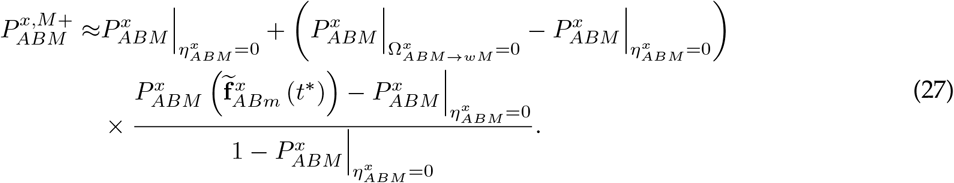

Here, 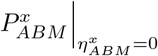 gives the probability of direct *ABM* establishment, while the parenthetic difference gives the additional probability attributable to an *ABm* lineage forming and establishing when paths through type-*wM* intermediates are excluded. The final factor, assuming that *ABM* lineages establishing under composition **f**_*R*_ (*t*^∗^) would also establish under 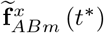, approximates the conditional probability that an *ABM* lineage establishes while the *ABm* lineage is becoming common given that direct establishment failed. Because this conditional probability is less than one, Eq. (27) reveals that 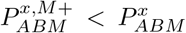 (Figure 2). This difference can arise because the modifier locus is unlinked to the major loci: segregation can separate *M* from the adaptive *AB* haplotype, leaving *M* associated with a deleterious genotype that may be lost through selection and genetic drift while the resulting *ABm* lineage establishes. Modifier alleles can therefore be short-lived, even when they induce establishment.

Despite this difference in magnitude, 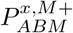 otherwise exhibits parameter dependence broadly similar to that of 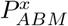: it grows with *δ*_*AB*_ (Figure 2A) and with reductions in *c* (Figure 2C) or *θ* (Figure 2D), with larger *θ* increasingly favoring recombination-rate over mutation-rate modification, 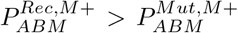. Like 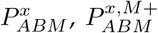 is only weakly affected by *δ*_*R*_ (Figure 2B), although it can increase slightly at larger values because—if *ABm* establishes—faster resident decline during the transition from resident to *ABm* commonality shifts the effective genotype-frequency composition toward *ABm* (Eq. (23)), increasing the replication rate of type-*ABM* organisms. This increase arises because an *ABm* mate already carries the *AB* haplotype, making offspring that inherit *M* more likely to also carry *AB*.

### 4.3 The probability of *wM* -induced establishment

Using an approach similar to that in the previous section, we now approximate the probability of *wM* - induced establishment, 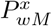. We account for direct paths, in which a type-*wM* lineage produces an *ABm* or *ABM* lineage that subsequently induces establishment, and paths through type-*vM* intermediates carrying haplotype *v* ≠*w*, where *v* ∈ {*Ab, aB*}. For simplicity, the latter paths are accounted for by treating type *vM* as a one-way path to establishment that does not produce new type-*wM* organisms, thereby excluding longer paths involving repeated transitions between *wM* and *vM* . Given this assumption, we first approximate the expected cumulative abundance of type-*vM* intermediates generated per type-*wM* organism, 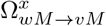. Substituting 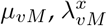, and 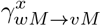 into Eq. (17) yields

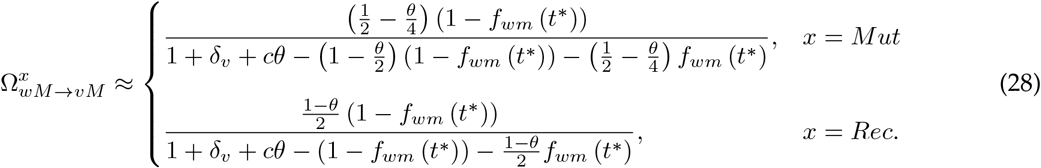

Each type-*vM* intermediate produces lineages that successfully induce establishment at a per-capita rate of 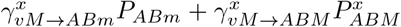. Multiplying this rate by 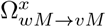 gives the contribution through type-*vM* intermediates. Type-*wM* organisms also contribute directly at a per-capita rate of 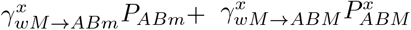. Summing these contributions and using the approximations of *P*_*ABm*_ and 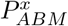 in Eqs. (19) and (22), respectively, gives the per-capita rate at which type-*wM* organisms produce lineages that successfully induce establishment:

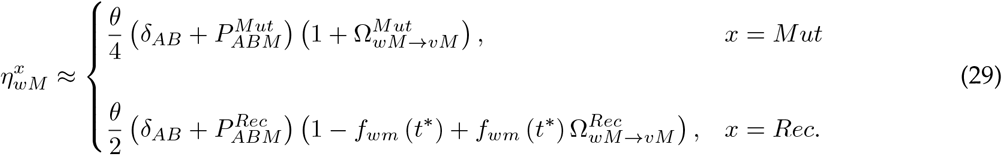

Thus, upon substituting 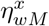 from Eq. (29)—along with *µ*_*wM*_ and 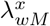 —into Eq. (16), we obtain

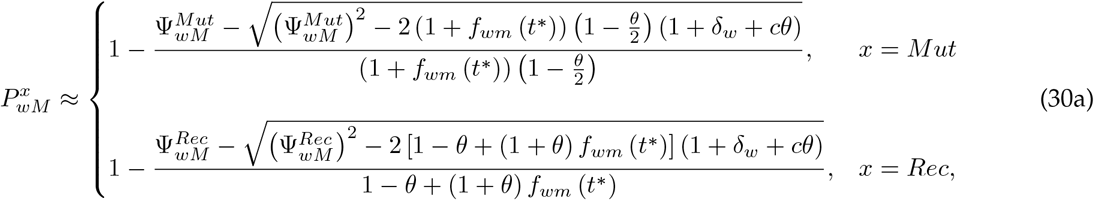

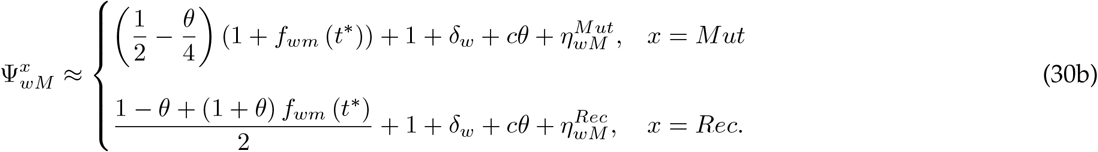

Equations (28)–(30) reveal a general tendency for reduced mortality among *M* -carrying organisms to promote *wM* -induced establishment, with Figure 3 illustrating these dependencies for a focal *AbM* lineage. Boosting *δ*_*AB*_ raises 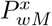 (Figure 3A) by increasing both *P*_*ABm*_ ≈ *δ*_*AB*_ and 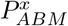, thereby augmenting the per-capita rate at which type-*wM* organisms produce lineages that induce establishment, 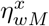, through the factor 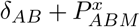 in Eq. (29). Whereas *δ*_*R*_ has only a weak-to-negligible effect on 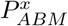, reducing *δ*_*R*_ substantially raises 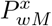 (Figure 3B) because the corresponding reduction in *δ*_*w*_ directly lowers the mortality rate of type-*wM* organisms rather than acting only through an intermediate-mediated path. Reducing *c* likewise raises 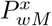 (Figure 3C) by diminishing *cθ*, which lowers the mortality rates of type-*wM* organisms and increases the expected cumulative abundance of type-*vM* intermediates, 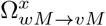.

**Figure 3:**
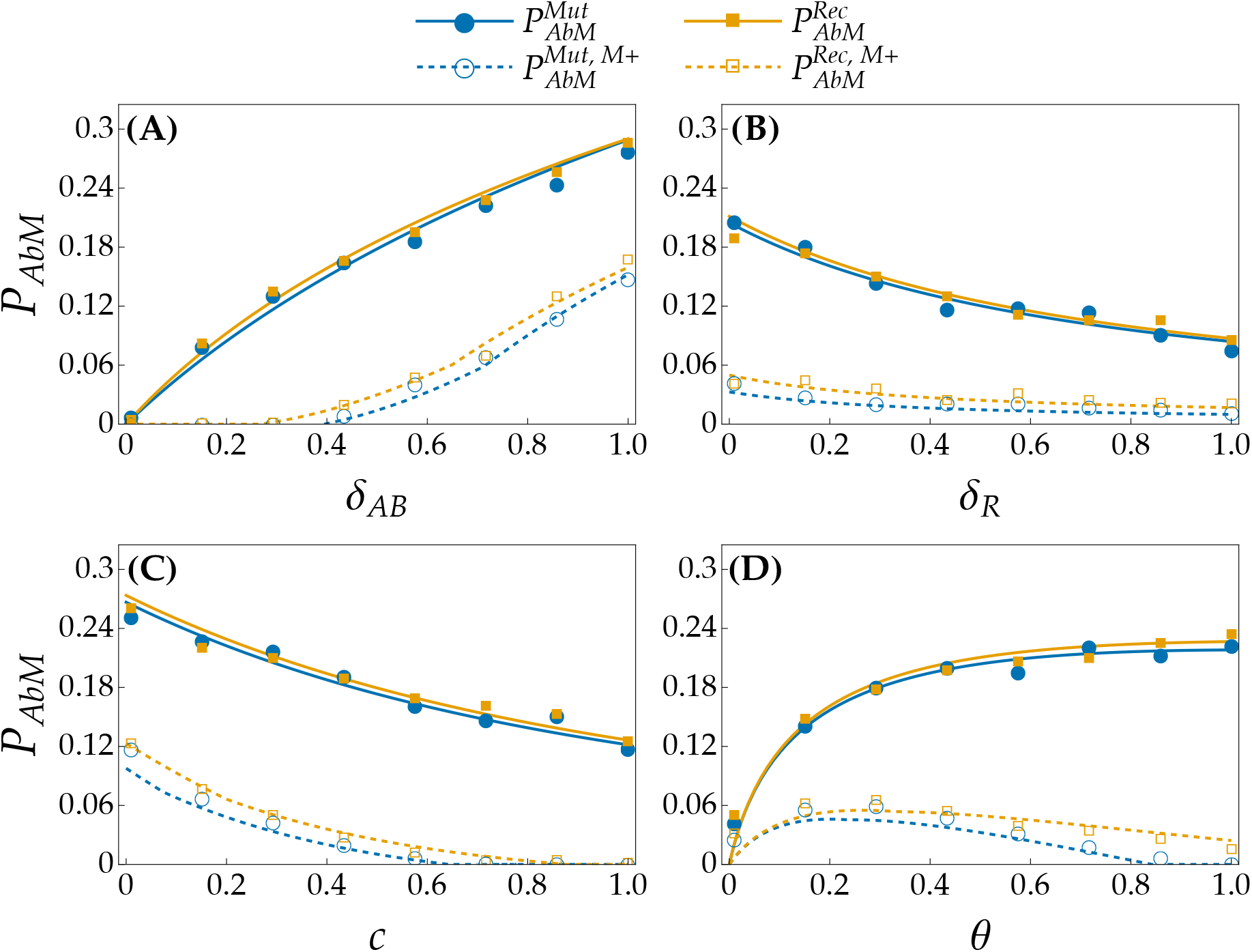
The probability of *AbM* -induced establishment, *P*_*AbM*_, and its constituent probability of *AbM* -induced establishment with modifier allele *M* persistence, 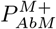, as functions of (A) the haplotype-*AB* mortality advantage, *δ*_*AB*_ ; (B) the resident mortality disadvantage, *δ*_*R*_; (C) the maximum mortality cost of *M* expression, *c*; and (D) the per-mating probability of modifier-induced genetic change, *θ*. Superscripts *Mut* and *Rec* denote the mutation- and recombinationrate modifier submodels, respectively. Each parameter affects the mortality rates of *M* -carrying organisms or their descendants, while *θ* additionally controls the rate of modifier-induced genetic change. Increasing *δ*_*AB*_ or reducing either *δ*_*R*_ or *c* raises *P*_*AbM*_ by lowering these mortality rates, whereas increasing *θ* boosts *P*_*AbM*_ by promoting production of descendants carrying the adaptive *AB* haplotype. As illustrated in each panel, the two modifier types can yield effectively equal *P*_*AbM*_ since they can produce *AB*-carrying descendants at similar rates, whereas recombination-rate modification can yield larger 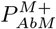 because of its tendency to better preserve *AB*. The behavior of 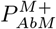 largely resembles that of *P*_*AbM*_, except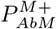 ‘s response to *θ* is nonmonotonic because a sufficiently large increase in *θ* augments the mortality cost of *M* expression enough to hinder *M* persistence. Since *M* can be lost after *ABM* forms, 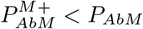. Markers: proportion of 3 *×* 10^3^ replicate simulations ending in establishment; solid curves: Eq. (30); dashed curves: Eq. (32). Unless stated otherwise, *c* = 0.3, *δ*_*AB*_ = 0.6, *δ*_*R*_ = 0.003, *δ*_*ab*_ = 10, *N*_0_ = 10^4^, *θ* = 0.5, *t*^∗^ = 1, and *U* = 0.

Unlike its effect on 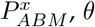 provides an exception to this mortality-driven pattern, with larger *θ* raising 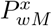 (Figure 3D) despite increasing *cθ* and, under mutation-rate modification, increasing the chance that mutation breaks apart the adaptive *AB* haplotype after it forms. This occurs because increasing *θ* also raises the rate at which type-*wM* organisms initially produce *ABm* or *ABM* descendants capable of inducing establishment. This effect is reflected in Eq. (29), where a boost in *θ* directly increases 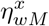 through the prefactors *θ/*4 and *θ/*2 under mutation- and recombination-rate modification, respectively. Under mutation-rate modification, the resulting increase in production of *AB*-carrying descendants outweighs both the greater mortality cost of *M* expression and the reduction in 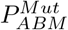 caused by mutation breaking apart *AB*. Although recombination-rate modification has the larger prefactor, *θ/*2, under our default regime of equal resident frequencies in **f**_*R*_ (*t*^∗^), *f*_*Abm*_ (*t*^∗^) = *f*_*aBm*_ (*t*^∗^) = 1*/*2, the frequency-dependent term under recombination-rate modification reduces its contribution to the same 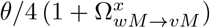 scaling as under mutation-rate modification. The two modifier types consequently tend to yield effectively equal probabilities of *wM* -induced establishment, 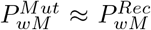 (Figure 3), when resident genotype frequencies are symmetric.

#### 4.3.1 The per-lineage probability of *wM* -induced establishment with modifier allele persistence

We next approximate the probability of *wM* -induced establishment with modifier allele *M* persistence, 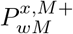. As with 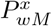, type-*vM* organisms are treated as one-way intermediates that do not produce new type-*wM* organisms. Since modifier allele persistence requires *ABM* establishment specifically, we exclude paths ending in *ABm* establishment while retaining both direct production of *ABM* by type-*wM* organisms and indirect production through type-*vM* intermediates. Using 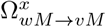 from Eq. (28), the direct percapita rate of *ABM* production is 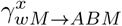while the corresponding rate through type-*vM* intermediates is approximately 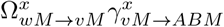. Summing these rates and multiplying by 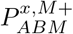 gives the per-capita rate at which type-*wM* organisms produce *ABM* lineages that establish with modifier allele persistence:

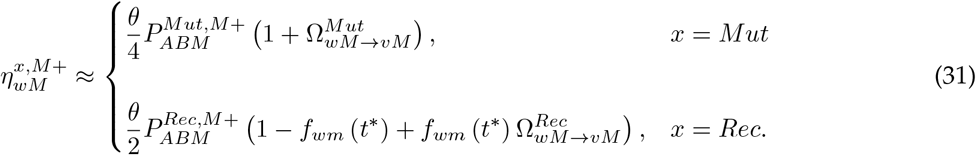

Substituting 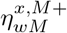 from Eq. (31)—as well as *µ*_*wM*_ and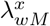 —into Eq. (16) and using Eq. (30) for 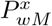 gives

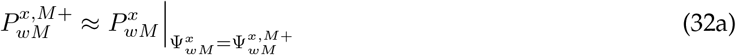

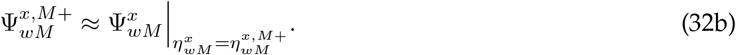

Since Eq. (32) is simply the expression for 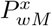 in Eq. (30) evaluated using 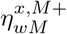 from Eq. (31), we see that, as with *ABM* -induced establishment, 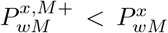 (Figure 3). This difference arises because establishment need not result in modifier allele persistence: whereas 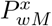 includes paths ending in either *ABm* or *ABM* establishment, 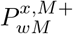 includes only paths producing an *ABM* lineage that establishes. Despite this difference in magnitude, 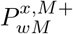 otherwise resembles 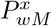 in its response to several model parameters: it increases with *δ*_*AB*_ (Figure 3A) and with reductions in *δ*_*R*_ (Figure 3B) or *c* (Figure 3C).

The effect of *θ* provides an important exception to this similarity: unlike 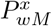, which increases monotonically with 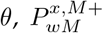 varies nonmonotonically with *θ* (Figure 3D). Starting from low *θ*, increasing *θ* raises 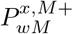 (Figure 3D) by boosting the rate at which type-*wM* organisms generate *ABM* lineages, as reflected by the factors *θ/*4 and *θ/*2 in Eq. (31). Once *θ* becomes sufficiently large, however, this benefit is outweighed by the growing *cθ*, causing 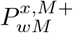 to decline with larger *θ* (Figure 3D). Because *M* persistence requires an *ABM* lineage to establish, paths through *ABm* descendants lacking *M* do not contribute to 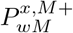 and hence cannot offset this decline. Relative to recombination-rate modification, the decline is more pronounced under mutation-rate modification (Figure 3D) because raising *θ* additionally increases the chance that mutation breaks apart *AB* after it forms, thereby further hindering *ABM* establishment. Consequently, under our default regime, recombination-rate modification tends to yield a larger probability of *wM* -induced establishment with modifier persistence, 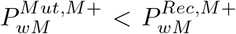 (Figure 3), particularly as *θ* becomes large.

### 4.4 The probability of ER

Having approximated the per-lineage probability of *wM* -induced establishment, 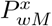, we can finally approximate the probability of ER, 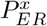. Modifier-allele-bearing lineages can arise throughout the resident decline, so the expected number that induce establishment, Λ^*x*^ (Eq. (14)), depends on both their rate of appearance and their establishment probabilities at their respective appearance times, *t*^∗^ = *t*_*M*_ . Demographic stochasticity causes realized resident genotype frequencies to vary among populations around those in **f**_*R*_ (*t*_*M*_ ) obtained from expected resident abundances, with stronger variation at smaller initial resident abundance, *N*_0_. Because 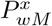 depends nonlinearly on resident genotype frequency, this variation can alter the mean per-lineage establishment probability averaged over appearance times and realized resident genotype frequencies, with each time-frequency state weighted by the rate of type-*wM* lineage appearance, which is proportional to the abundance of type *wm*. Under our default regime of *δ*_*Ab*_ = *δ*_*aB*_ = *δ*_*R*_, for which the frequencies in **f**_*R*_ (*t*_*M*_ ) obtained from expected resident abundances remain constant during the decline, we account analytically for these effects using the diffusion-approximated type-*wm*-weighted mean, 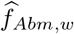, and variance, *V*_*w*_, in realized type-*Abm* frequency under this appearance-weighted distribution. The former determines the corresponding type-*wm*-weighted mean resident genotype-frequency composition,

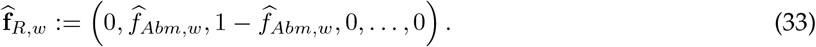

Because resident genotype-frequency composition is determined by the frequency of type *Abm*, a secondorder expansion of 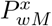 around 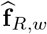 then gives the appearance-weighted mean per-lineage establishment probability adjusted for stochastic resident genotype-frequency variation (Appendix F),

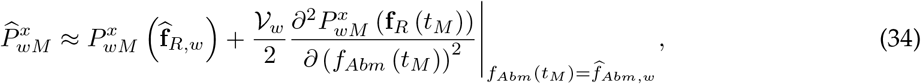

which we combine with the cumulative expected number of type-*wM* lineage appearances over the resident decline to obtain their contribution to Λ^*x*^. When *δ*_*Ab*_ *> δ*_*aB*_, stochastic variation in resident genotype frequency likewise occurs but, for simplicity, we do not account for it analytically. In this regime, type *Abm* declines more rapidly than type *aBm*, so the resident composition, **f**_*R*_ (*t*_*M*_ ), obtained from expected resident abundances changes through time, as does 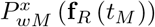. Rather than derive a corresponding adjustment for stochastic resident genotype-frequency variation, we therefore evaluate 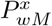 using **f**_*R*_ (*t*_*M*_ ) and approximate this temporal change over the decline. To retain a closed-form approximation of Λ^*x*^, we represent the change using two phases separated by a transition time, *t*_*c*_. The establishment probability associated with the initial resident composition is used for 0 ≤ *t*_*M*_ *< t*_*c*_, whereas that associated with the limiting composition is used for *t*_*M*_ ≥ *t*_*c*_. We define *t*_*c*_ as the time at which the type-*Abm* frequency in**f**_*R*_ (*t*_*M*_ ) falls to half its initial value, *f*_*Abm*_ (*t*_*c*_) = *f*_*Abm*_ (0) */*2, obtaining

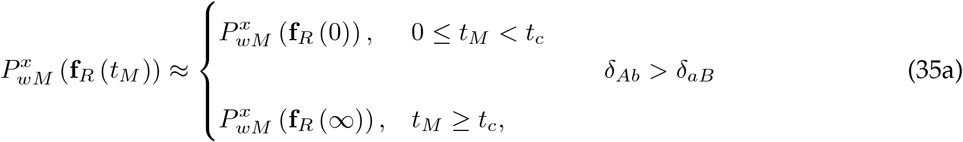

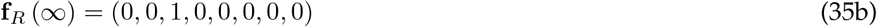

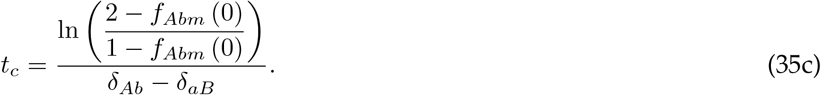

Our treatments of the equal- and unequal-resident-mortality-disadvantage regimes therefore require different evaluations of Λ^*x*^ in Eq. (14a). Under equal resident mortality disadvantages, integrating the type-*AbM* and type-*aBM* lineage-appearance rates in Eq. (14a) over the resident decline yields cumulative expected numbers of lineage appearances of *Uf*_*Abm*_(0)*/δ*_*R*_ and *U* (1 − *f*_*Abm*_(0))*/δ*_*R*_, respectively. Multiplying each of these quantities by its corresponding appearance-weighted mean per-lineage establishment probability in Eq. (34) gives 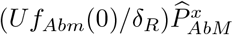 for type-*AbM* and 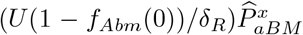 for type-*aBM* . Summing these two expected numbers of lineage appearances that induce establishment yields the equal-disadvantage approximation of Λ^*x*^. Under unequal resident mortality disadvantages, we instead substitute the piecewise establishment probabilities in Eq. (35) into Eq. (14a) and integrate over 0 ≤ *t*_*M*_ *< t*_*c*_ and *t*_*M*_ ≥ *t*_*c*_. Substituting the resulting approximations of Λ^*x*^ into Eq. (12) yields

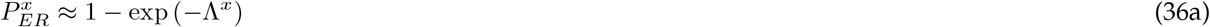

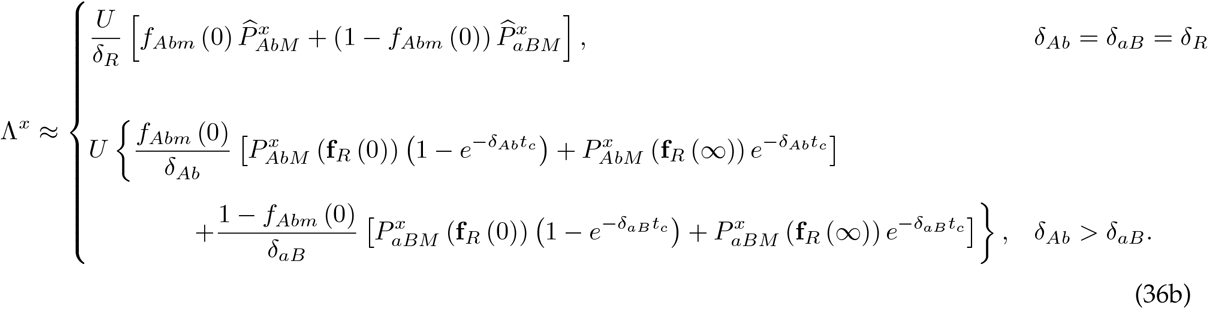

Much of the qualitative behavior of 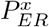 resembles that of 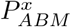 (Figures 3–4), with lower mortality rates among *M* -carrying organisms and their descendants tending to promote ER (Figure 4). Increasing *δ*_*AB*_ raises 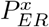 (Figure 4A) by augmenting the per-lineage probabilities of establishment across all *M* appearance times, *t*_*M*_, whereas reducing *δ*_*R*_ boosts 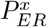 (Figure 4B) both by increasing these probabilities and, more importantly, by increasing the time to resident extinction and hence the cumulative opportunity for modifier-locus mutation across all *t*_*M*_, as reflected by the factor 1*/δ*_*R*_ in Eq. (36b). Similarly, reducing *c* enhances 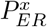 (Figure 4C) by decreasing *cθ*, whereas increasing *θ* boosts 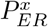 (Figure 4D) by raising the rate at which *M* -bearing lineages produce *AB* descendants that can establish despite the accompanying increase in *cθ*.

**Figure 4:**
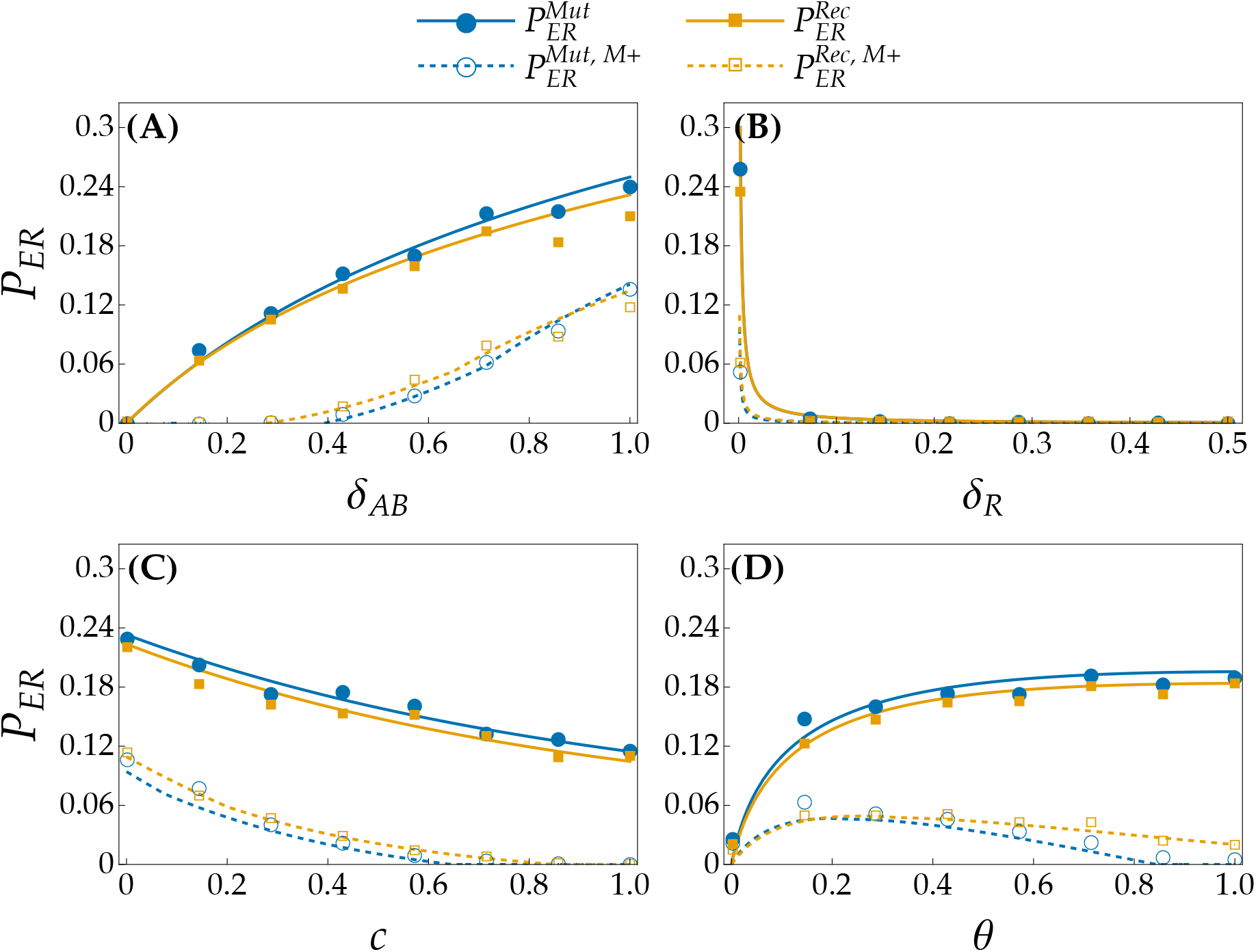
The probability of ER, *P*_*ER*_, and its constituent probability of ER with modifier allele *M* persistence, 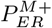, as functions of (A) the haplotype-*AB* mortality advantage, *δ*_*AB*_ ; (B) the resident mortality disadvantage, *δ*_*R*_; (C) the maximum mortality cost of *M* expression, *c*; and (D) the per-mating probability of modifier-induced genetic change, *θ*. Superscripts *Mut* and *Rec* denote the mutation- and recombination-rate modifier submodels, respectively. Each parameter affects the mortality rates of *M* -carrying organisms or their descendants, while *θ* additionally controls the rate of modifier-induced genetic change. Increasing *δ*_*AB*_ or reducing either *δ*_*R*_ or *c* raises *P*_*ER*_ by lessening these mortality rates, with lower *δ*_*R*_ additionally prolonging the resident decline and thereby increasing the opportunity for modifier-locus mutation. Increasing *θ* instead raises *P*_*ER*_ by promoting production of descendants carrying the adaptive *AB* haplotype despite augmenting the mortality cost of *M* expression. Panels (A), (C), and (D) show that mutation-rate evolution can yield larger *P*_*ER*_ under some conditions, despite recombination-rate evolution tending to yield larger 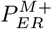 due to its tendency to better preserve *AB*. The behavior of 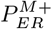 broadly resembles that of *P*_*ER*_ but responds nonmonotonically to *θ* because a sufficiently large *θ* increases the mortality cost of *M* expression enough to hinder *M* persistence. Since *M* can be lost during *AbM* -induced establishment, 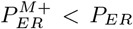. Markers: proportion of 3 *×* 10^3^ replicate simulations ending in ER; solid curves: Eq. (36); dashed curves: Eq. (37). Unless stated otherwise, *c* = 0.3, *δ*_*AB*_ = 0.6, *δ*_*R*_ = 0.003, *δ*_*ab*_ = 10, *N*_0_ = 10^4^, *θ* = 0.5, and *U* = 0.003.

Under our default regime, for which *f*_*Abm*_ (*t*_*M*_ ) = *f*_*aBm*_ (*t*_*M*_ ) = 1*/*2 in **f**_*R*_ (*t*_*M*_ ) at every *t*_*M*_, the probability of ER via mutation-rate evolution, 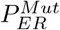, and that via recombination-rate evolution, 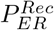, can be effectively equal when either *δ*_*AB*_ is sufficiently small (Figure 4A) or *θ* is sufficiently small (Figure 4D), paralleling the near-equivalence of 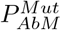 and 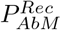 in Figure 3. Reducing *δ*_*AB*_ weakens the establishment advantage of *AB*-carrying descendants and therefore the consequences of modifier-specific differences after *AB* forms, whereas decreasing *θ* makes the two modifier types generate *AB* at effectively equal rates under symmetric resident frequencies and reduce the consequences of their differences in modifier-induced genetic change. Outside this near-equivalence, however, 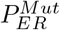 can exceed 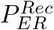 (Figure 4D), and whether this occurs also depends on the initial resident abundance, *N*_0_ (Figure 5A). Decreasing *N*_0_ strengthens demographic stochasticity, causing realized resident frequencies at a given *t*_*M*_ to fluctuate more strongly around 1*/*2. When one resident genotype is stochastically more common, *M* -bearing lineages arise more frequently from that genotype, but the complementary resident haplotype required for those lineages to generate *AB* by recombination is correspondingly rarer. Mutation does not require this complementary haplotype and is therefore less sensitive to such frequency fluctuations. Consequently, demographic stochasticity lowers the appearance-weighted mean per-lineage establishment probabilities, 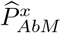 and 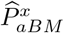 (Eq. (34)), more strongly under recombination-rate modification, allowing 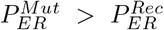 (Figure 5A) even when 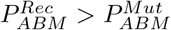 (Figure 2).

**Figure 5:**
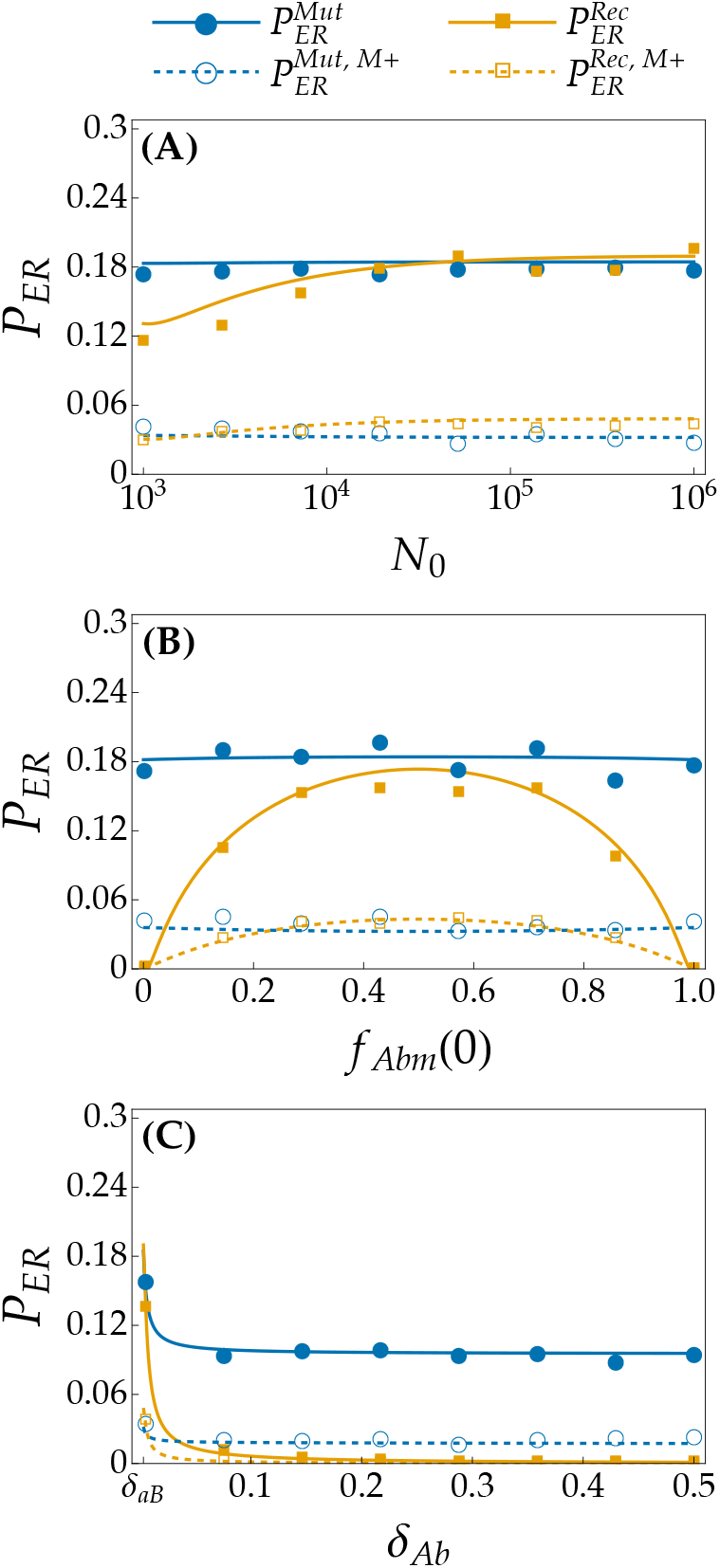
The probability of ER, *P*_*ER*_, and its constituent probability of ER with modifier allele *M* persistence, 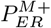, as functions of (A) the initial resident abundance, *N*_0_; (B) the initial type-*Abm* frequency, *f*_*Abm*_(0); and (C) the type-*Abm* mortality disadvantage, *δ*_*Ab*_, with the haplotype-*aB* disadvantage, *δ*_*aB*_, held fixed. Superscripts *Mut* and *Rec* denote the mutation- and recombination-rate modifier submodels, respectively. Changes to each parameter can induce asymmetry between the realized resident genotype frequencies: reducing *N*_0_ strengthens demographic stochasticity and hence stochastic frequency asymmetry; moving *f*_*Abm*_(0) away from 1*/*2 deterministically makes one resident genotype initially more common than the other; and increasing *δ*_*Ab*_ above *δ*_*aB*_ causes type *Abm* to decline more rapidly on average than type *aBm*. Each panel reveals that greater resident-frequency asymmetry tends to favor mutation-rate evolution for *P*_*ER*_ because recombination is increasingly constrained by the rarer complementary resident haplotype required to generate *AB*. With sufficient symmetry, however, the two routes to ER can yield effectively equal *P*_*ER*_. The behavior of 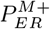 broadly resembles that of *P*_*ER*_, except that sufficient symmetry can instead yield 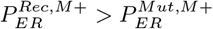 because of recombination-rate modification’s tendency to better preserve *AB*. Markers: proportion of 3 *×* 10^3^ replicate simulations ending in ER; solid curves: Eq. (36); dashed curves: Eq. (37). Unless stated otherwise, *c* = 0.3, *δ*_*AB*_ = 0.6, *δ*_*Ab*_ = *δ*_*aB*_ = *δ*_*R*_ = 0.003, *f*_*Abm*_(0) = 1*/*2, *N*_0_ = 10^4^, *θ* = 0.5, and *U* = 0.003.

Resident-frequency asymmetry arising deterministically can likewise favor mutation-rate evolution. One source is *f*_*Abm*_(0) ≠*f*_*aBm*_(0) (Figure 5B), which shifts the relative contributions of the two *M* -bearing lineage paths in Eq. (36b) through the weights *f*_*Abm*_(0) and *f*_*aBm*_(0). Under mutation-rate evolution, decreasing either resident frequency necessarily increases the other, so the reduced contribution of one symmetric mutation-mediated path is largely offset by the increased contribution of the other, leaving 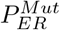 comparatively unchanged. Under recombination-rate evolution, greater asymmetry instead makes the complementary resident haplotype required to generate *AB* increasingly rare, further reducing 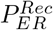 relative to 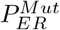 (Figure 5B).

Another form of deterministic resident-frequency asymmetry that can favor mutation-rate evolution arises when *δ*_*Ab*_ is increased independently of *δ*_*aB*_ (Figure 5C). With *δ*_*aB*_ held fixed, increasing *δ*_*Ab*_ causes type *Abm* to decline increasingly rapidly on average, systematically shifting the resident population toward type *aBm*. Mutation-rate evolution nevertheless retains a route to ER because type *aBM* can generate *AB* by mutation without type *Abm*, such that 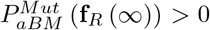 in Eq. (36b) and 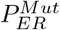 approaches a nonzero limit. Under recombination-rate evolution, by contrast, 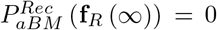 because the complementary type-*Abm* haplotype is absent, causing 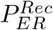 to approach zero (Figure 5C).

### 4.5 The probability of ER with modifier allele persistence

We next approximate the probability of ER with modifier allele persistence, 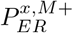. Replacing the perlineage establishment probabilities in Λ^*x*^ with their counterparts requiring modifier allele persistence gives

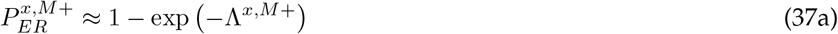

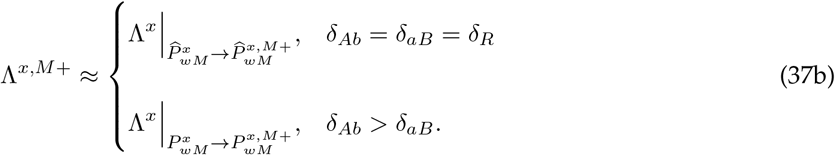

Just as 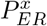 qualitatively resembles 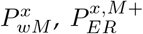 largely mirrors 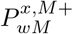 (Figures 3–4). In particular, 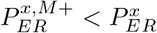 (Figure 4) because ER need not preserve the modifier allele driving it. Increasing *δ*_*AB*_ raises 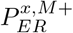 (Figure 4A), whereas augmenting *δ*_*R*_ or *c* lowers it (Figure 4B–C). Like 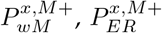 also varies nonmonotonically with *θ* (Figure 4D): raising *θ* initially promotes production of *M* -bearing *AB* descendants, whereas a sufficiently large *θ* enhances *cθ* enough to hinder modifier persistence. Notably, even when mutation-rate evolution yields a higher probability of ER, recombination-rate evolution can yield a higher probability of ER with modifier persistence, 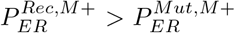 (Figure 4D), because recombination better preserves *AB* after it forms.

Whether this advantage of recombination-rate evolution is notable depends largely on *N*_0_ (Figure 5A). At smaller *N*_0_, the stochastic frequency asymmetry that favors mutation-rate evolution for 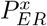 produces only a weak corresponding advantage in 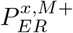, such that 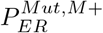 and 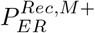 can remain effectively equivalent (Figure 5A). As increasing *N*_0_ weakens these fluctuations, however, recombination-rate modification’s tendency to better preserve *AB* can eventually cause 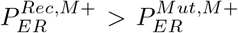 (Figure 5A). In contrast, deterministic resident-frequency asymmetry favors mutation-rate evolution for both 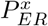 and 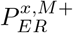: changing *f*_*Abm*_(0) away from 1*/*2 (Figure 5B) or raising *δ*_*Ab*_ independently of *δ*_*aB*_ (Figure 5C) increasingly favors mutation-rate evolution. In the latter case, increasing *δ*_*Ab*_ causes 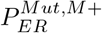 to approach a nonzero limit because type *aBM* can generate *ABM* without type *Abm*, whereas 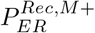 approaches zero because recombination requires the increasingly rare complementary type *Abm*.

## Discussion

Here, we analytically approximated and compared the probability of ER through two alternative routes— mutation- and recombination-rate evolution—along with the probability that the modifier allele *M* facilitating ER persists through rescue. We considered two complementary resident genotypes, *Abm* and *aBm*, from which either mutation or recombination can produce the same adaptive haplotype, *AB*, carried as type *ABm* without *M* or as type *ABM* with it. Our results show that sufficiently strong asymmetry between resident genotype frequencies can favor mutation-rate evolution (Figure 5), whether arising through stochastic frequency fluctuations, unequal initial frequencies, or more rapid loss of one genotype. This occurs because mutation can generate *AB* from either resident genotype, whereas recombination becomes increasingly constrained by the scarcity of complementary haplotypes. Under sufficiently symmetric resident frequencies, by contrast, the two routes can yield effectively equal probabilities of ER (Figure 5). Even when ER through either route is appreciably likely, however, the responsible modifier allele is often lost before rescue is complete, such that the probability of ER with *M* persistence remains lower than the overall probability of ER (Figure 4). Multiple factors contribute to this difference. Segregation can separate *M* from *AB*, allowing a type-*ABm* lineage to establish after the modifier allele is lost. The fitness cost of *M* expression can further hinder modifier persistence, with the probability of *M* persistence peaking at an intermediate probability of modifier-induced genetic change (Figure 3) due to the fitness cost becoming too large relative to the benefit of *AB* production. Continued modifier-induced mutation can also break apart *AB* after it forms (Figure 2). Since recombination between *AB* and either resident haplotype cannot disrupt *AB*, this mutation-specific effect can make recombination-rate modifiers more likely to persist than mutation-rate modifiers even when mutation-rate evolution yields the higher probability of ER (Figure 4).

### 4.6 The relative effectiveness of mutation- and recombination-rate evolution as ER mechanisms can depend less on their intrinsic differences than on resident genotype composition

A substantial body of theoretical and empirical work has examined the evolution of mutation rates (Gifford et al., 2023; Ishii et al., 1989; Liberman & Feldman, 1986; Liberman et al., 2011) and recombination rates (Aggarwal et al., 2015; Barton, 1995; Charlesworth, 1976; Dapper & Lively, 2014; Feldman et al., 1980; Otto & Michalakis, 1998). Among theoretical studies, many have considered evolution over relatively long timescales in populations of effectively infinite size (e.g., Feldman et al., 1980; Ishii et al., 1989; Liberman & Feldman, 1986), rather than the short-term evolution of finite, declining populations characteristic of ER. Especially relevant is the work of Carja et al. (2014), who compared evolutionarily stable mutation and recombination rates in infinite haploid populations experiencing changing selective environments. For periodic environmental change, they considered alternate environments that favored different genetic states with equal strength, preventing a systematic advantage of either state, while infinite population size eliminated stochastic departures from this symmetry (Carja et al., 2014). Under these conditions, the evolutionarily stable mutation and recombination rates responded qualitatively similarly, with both declining as the interval between environmental shifts increased because the benefit of generating new genetic variation diminished during longer periods of constant selection (Carja et al., 2014). Our work therefore complements theirs by identifying a different dimension along which mutation- and recombination-rate evolution can be similar: following a single environmental shift, the two can be comparably successful routes to ER when the complementary resident genotypes required for recombination to generate *AB* remain sufficiently symmetric in frequency. As this symmetry is lost, however, mutation-rate evolution becomes increasingly favored because mutation can generate *AB* from either resident genotype independently, whereas recombination becomes constrained by the rarer complementary genotype. Thus, the relative effectiveness of the two mechanisms can be determined less by their intrinsic differences in generating adaptive variation than by the resident genotype composition on which they act.

### 4.7 Modifier alleles need not persist to facilitate ER

Closer to the demographic setting considered here are the relatively few studies of mutation-rate evolution during ER (Greenspoon & Mideo, 2017; Tuffaha & Wahl, 2025). Greenspoon and Mideo (2017), in particular, considered mutation-rate evolution during ER by simulating parasite populations under cycling drug treatment. They showed that a modifier allele conferring an elevated mutation rate could increase in frequency through its association with newly favored resistance alleles, whereas recombination between the modifier and resistance loci weakened this association and reduced the extent to which mutation-rate evolution facilitated ER (Greenspoon & Mideo, 2017). Complementing this result, our analysis shows that, when the modifier locus is unlinked, segregation can separate *M* from *AB* under either form of rate evolution, allowing an *ABm* lineage to establish while reducing the probability of ER with *M* persistence. Experimental results from Raynes et al. (2011) support this prediction by showing that an allele conferring an elevated mutation rate increased in asexual diploid yeast populations but declined toward loss in sexual diploid populations (Raynes et al., 2011). Although the authors found that the allele’s realized fitness disadvantage was sufficient to explain this decline, they noted that recombination could additionally separate the elevated-mutation-rate allele from beneficial mutations (Raynes et al., 2011). Together with this empirical evidence for the underlying mechanism, our results reveal that modifier alleles need not persist to facilitate ER.

### 4.8 The form of rate evolution most likely to facilitate ER need not leave the most persistent modifier allele

In another study of mutation-rate evolution during ER, Tuffaha and Wahl (2025) used numerical approximations validated by simulations to estimate ER probabilities in a model with complete linkage between a mutation-rate modifier locus and a single major locus (Tuffaha & Wahl, 2025). In their model, a modifier allele confers a genome-wide mutation rate *F* -fold higher than that of the wild type, thereby increasing the rate at which an adaptive allele arises, while back mutation is ignored (Tuffaha & Wahl, 2025). They found that ER probability can peak at intermediate wild-type mutation rates as increasing mutational input is eventually offset by mutation-associated fitness costs (Tuffaha & Wahl, 2025). Our analytical approximations both recover and extend this qualitative result: under mutation-rate evolution, the probability of ER with *M* persistence can likewise peak at an intermediate probability of modifier-induced genetic change because sufficiently high probabilities not only increase the mortality cost of *M* expression but, when back mutation is permitted, also raise the chance that mutation disrupts *AB* after it forms. This additional cost of continued mutation can make mutation-rate modifiers less likely to persist than recombination-rate modifiers because recombination tends to preserve *AB* in descendants of type *ABM*, whereas mutation can continue to break the adaptive haplotype apart. Therefore, the form of rate evolution that most strongly promotes rescue can differ from the one whose modifier is most likely to remain in the rescued population.

### 4.9 Model assumptions and future directions

Our model is built on several simplifying assumptions that could be relaxed in future work. A major set concerns the genetic architecture of modifier action and the association between the modifier locus and the adaptive haplotype. We assumed that the resident allele at the modifier locus confers negligible background rates of mutation and recombination at the major loci, consistent with the reduction principle whereby selection near equilibrium tends to favor alleles that reduce rates of variation production (Altenberg et al., 2017; Feldman & Liberman, 1986). Allowing non-negligible background rates could reveal a threshold rate of major-locus genetic change above which ER is more likely than not without the appearance of the modifier allele. We also assumed that the modifier locus is unlinked to the major loci; tighter linkage could both promote modifier allele persistence by preventing the allele’s segregation away from the adaptive haplotype and make modifier-locus position consequential for whether newly generated adaptive genotypes retain the modifier allele. Finally, we assumed that the modifier allele does not alter mutation at its own locus or recombination involving that locus. Relaxing this assumption could introduce feedback between modifier evolution and persistence, potentially allowing an ER-facilitating modifier to promote its own loss from the resulting adaptive lineage.

A separate assumption concerns the genetic composition of the resident population. We assumed that the *ab* haplotype is sufficiently deleterious to be absent from the resident population and immediately purged if produced during population decline. Our model is therefore effectively equivalent to one in which *ab* is the predominant wild-type haplotype before environmental change, with *Ab* and *aB* present as standing genetic variants, but is eliminated immediately after the change, leaving only *Ab* and *aB*. In an analysis of ER under a fixed recombination rate, Uecker and Hermisson (2016) showed that when wild-type *ab* instead remains common and declines slowly, *Ab* and *aB* are less likely to encounter one another and recombine, while any *AB* formed is more likely to be broken apart through recombination with *ab* (Uecker & Hermisson, 2016). By isolating the case in which these complications are absent, our model provides a baseline understanding of when recombination-rate evolution can provide a viable route to ER. Extending the model to allow *ab* to persist could identify a threshold initial frequency or rate of *ab* decline at which mutation-rate evolution becomes a more successful route to ER than recombination-rate evolution.

Finally, the derivation of the ER probability entailed its own set of simplifying assumptions. We treated each focal lineage arising during ER as rare and the surrounding genotype-frequency composition as effectively constant during its initial stochastic dynamics, permitting branching-process approximations of per-lineage establishment probabilities. To account for ER paths passing through transient intermediates carrying the resident major-locus haplotypes *Ab* or *aB* before forming or reforming *AB*, we approximated the expected cumulative abundance of each intermediate and its resulting contribution to *AB* production. To capture cases in which *ABM* establishes after *ABm*, we used an effective genotype-frequency composition as a proxy for the changing composition as *ABm*’s frequency rises and the resident genotype frequencies decline. Under equal resident mortality disadvantages, we further used diffusion theory to account for stochastic resident-frequency fluctuations. Even with these corrections, however, the resulting ER probability approximation loses quantitative accuracy once initial resident abundance becomes sufficiently small and these fluctuations sufficiently strong (Figure 5A). A challenge for future theory is to develop analytical approaches that better capture both the time-varying expected genotype-frequency composition and stochastic fluctuations around it while remaining interpretable. Such approaches could permit tractable ER approximations for more genetically and ecologically complex scenarios that, when compared, yield simple rules of thumb for when populations are most likely to persist by evolving their own capacity to generate adaptive variation.

## Study Funding

This work was supported by U.S. National Science Foundation grant 2143063 awarded to A.L.D.

## Appendix A Coefficients comprising the mutation-rate modifier submodel

Conditional on a major-locus mutation, the parental gamete receiving the mutation and the affected major locus are each chosen at random, while the segregation of haplotype *i* with allele *M* and the offspring assigned to the focal female population are likewise determined at random. In each case, the choice is between two equally likely alternatives, with either alternative occurring with probability 1*/*2. Accordingly, each coefficient is obtained by multiplying the probabilities of the independent choices required for the focal outcome; a factor of 1*/*2 appears only when one of the two alternatives is required and is omitted when either alternative produces the same outcome. For an *M* × *m* mating with major-locus Hamming distance *H*(*i, j*) = 0, both parental haplotypes are *i*, so no mutation preserves *i* regardless of modifier-locus segregation, whereas a mutation leaves only one parental copy of *i* available for replication:

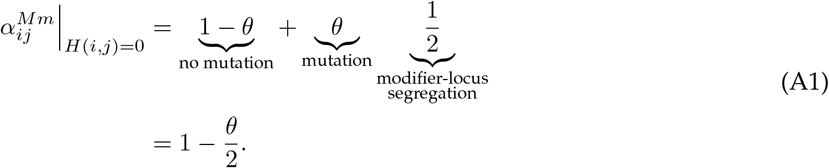

When *H*(*i, j*) = 1, mutation can leave the existing copy of *i* available for replication and, when the differing locus of *j* is mutated, additionally generate *i* from *j*:

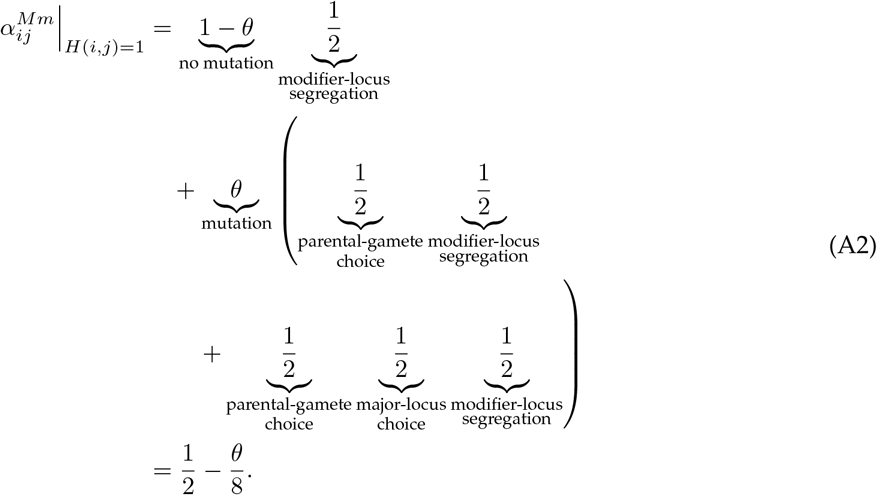

When *H*(*i, j*) = 2, a single mutation cannot generate *i* from *j*, so replication following mutation requires the existing copy of *i* to be retained:

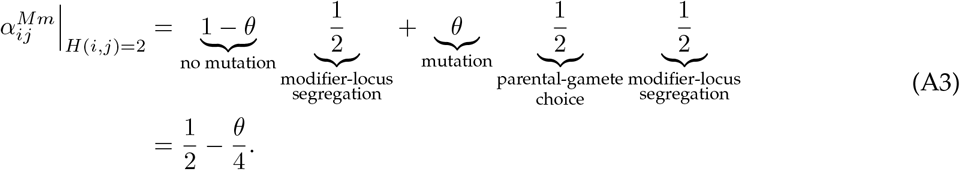

For *M* × *M* matings, the same mutational paths apply, except that modifier-locus segregation is absent. When *H*(*i, j*) = 0, both parental haplotypes are *i*, and a mutation leaves one copy of *i* whose assignment to the focal female population occurs with probability 1*/*2:

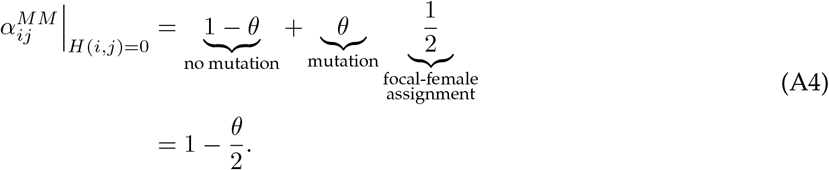

When *H*(*i, j*) = 1, mutation can again leave the existing copy of *i* available for replication and, when the differing locus of *j* is mutated, additionally generate *i* from *j*:

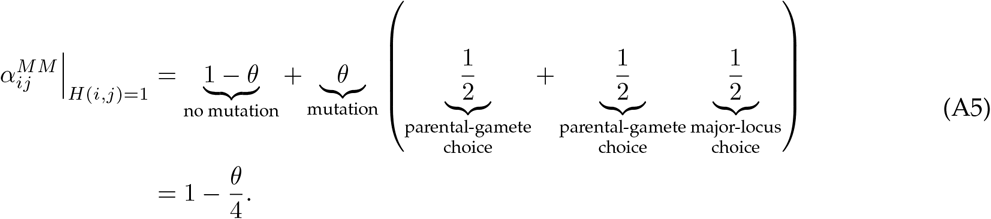

When *H*(*i, j*) = 2, a single mutation cannot generate *i* from *j*, leaving only retention of the existing copy of *i*:

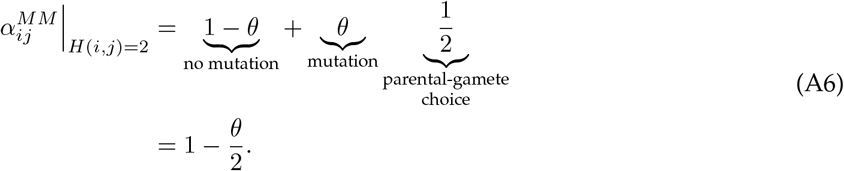

For genetic input, 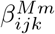 gives the per-mating-direction probability of generating the focal haplotype *i* from nonfocal parental haplotypes *j* and *k*. When *H*(*i, j*) + *H*(*i, k*) = 2, both parental haplotypes are one mutation from *i*, so either parental gamete can be chosen and only the correct major locus must be mutated:

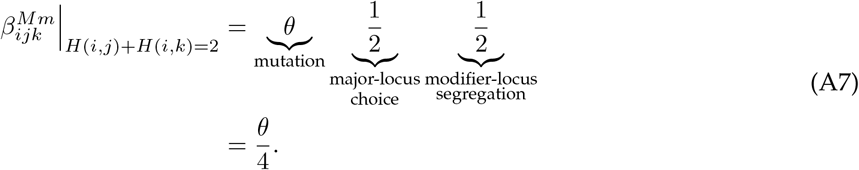

When *H*(*i, j*) + *H*(*i, k*) = 3, only one parental haplotype is one mutation from *i*, so both that parental gamete and its differing major locus must be chosen:

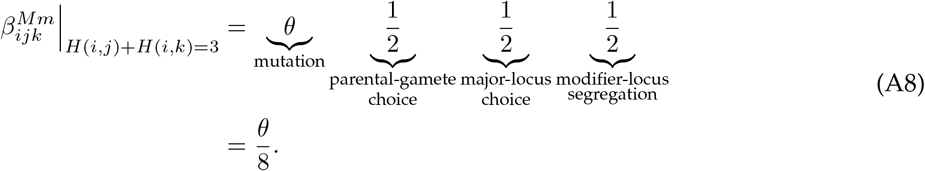

When *H*(*i, j*) + *H*(*i, k*) = 4, neither parental haplotype can generate *i* by a single mutation:

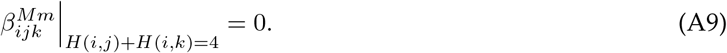

Removing the 1*/*2 modifier-locus segregation factor in an *M* × *M* mating gives the corresponding genetic-input probability, 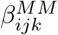:

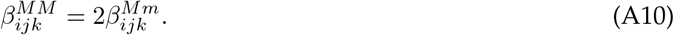

## Appendix B Coefficients comprising the recombination-rate modifier submodel

Recombination can alter a major-locus haplotype only when the parental haplotypes differ at both major loci. Thus, unlike mutation, no additional choice of parental gamete or major locus is involved: a crossover either changes the parental haplotypes when they differ at both major loci or leaves them unchanged when they do not. For an *M* × *m* mating with *H*(*i, j*) = 0, both parental haplotypes are *i*, so recombination cannot alter the focal haplotype:

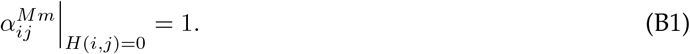

When *H*(*i, j*) = 1, recombination likewise cannot alter the parental haplotypes, leaving modifier-locus segregation as the only constraint:

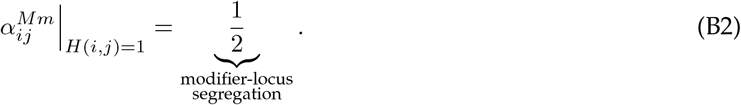

When *H*(*i, j*) = 2, replication additionally requires that no crossover occur:

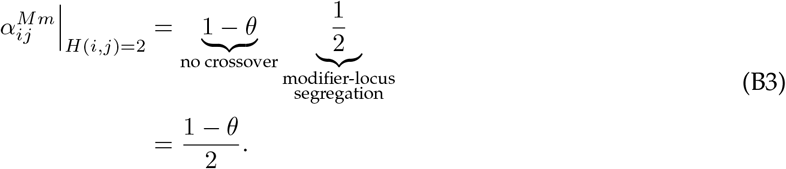

For *M* ×*M* matings, modifier-locus segregation is absent, so recombination affects replication only when the parental haplotypes differ at both major loci:

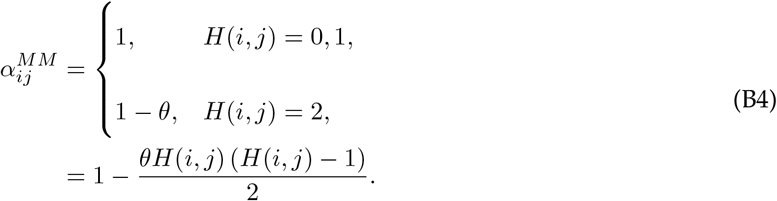

For genetic input from nonfocal haplotypes *j, k* ≠*i*, recombination can generate *i* only when *H*(*j, k*) = 2 and *H*(*i, j*) + *H*(*i, k*) = 2. Under these conditions, a crossover generates the focal recombinant haplotype, leaving modifier-locus segregation as the remaining constraint:

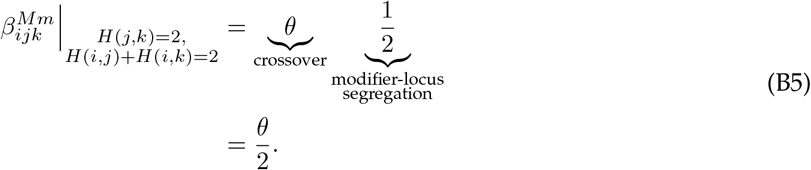

Otherwise,

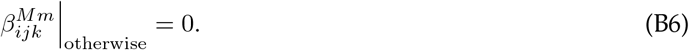

Removing the 1*/*2 modifier-locus segregation factor in an *M* × *M* mating doubles the genetic-input probability:

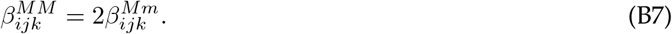

## Appendix C The per-lineage probability of *h*-induced establishment

To approximate the per-lineage probability of *h*-induced establishment, *P*_*h*_ (**f** (*t*^∗^)), we apply branching process theory (Stana et al., 2025; Uecker et al., 2015), assuming that nonfocal genotype frequencies remain effectively constant prior to establishment and that type-*h* organisms remain sufficiently rare for their frequency and modifier-locus mutation rate to be negligible. Consequently, descendant lineages founded by different type-*h* organisms are independent and experience the same per-capita rates at **f** (*t*^∗^). For any ℓ ∈ *L*_*h*_, type *h* produces type-ℓ lineages at per-capita rate *γ*_*h*→ℓ_ (**f** (*t*^∗^)), and each such lineage induces establishment with probability *P*_ℓ_ (**f** (*t*^∗^)). Summing across *L*_*h*_ therefore gives the total per-capita rate at which type *h* produces lineages that successfully induce establishment,

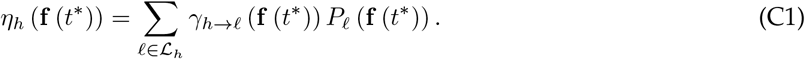

The combined per-capita rate of type-*h* replication, death, or production of a lineage that successfully induces establishment is then

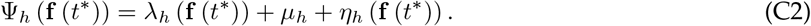

We define *ρ*_*h*_ (**f** (*t*^∗^)) = 1 − *P*_*h*_ (**f** (*t*^∗^)) as the probability that a lineage founded by one type-*h* organism does not induce establishment. Over the next infinitesimal interval (*t, t* + *dt*), none of the processes contributing to Ψ_*h*_ occurs with probability 1 − Ψ_*h*_ (**f** (*t*^∗^)) *dt* + *O dt*^2^ ; replication occurs with probability *λ*_*h*_ (**f** (*t*^∗^)) *dt*, after which the parental organism and its offspring independently avoid establishment with probability 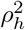; death occurs with probability *µ*_*h*_*dt* and guarantees non-establishment; and production of a lineage that successfully induces establishment guarantees establishment. Thus,

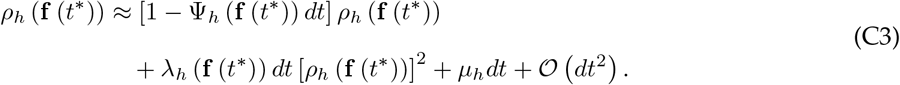

Subtracting *ρ*_*h*_ (**f** (*t*^∗^)), dividing by *dt*, and taking *dt* → 0 gives

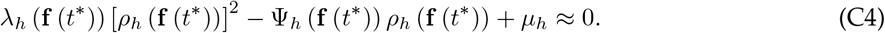

Taking the minimal root in [0, 1] and substituting *P*_*h*_ (**f** (*t*^∗^)) = 1 − *ρ*_*h*_ (**f** (*t*^∗^)) yields

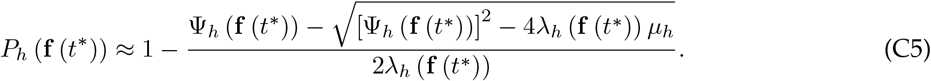

## Appendix D Table of per-capita rates used in the analytical approximations

**Table D1:**
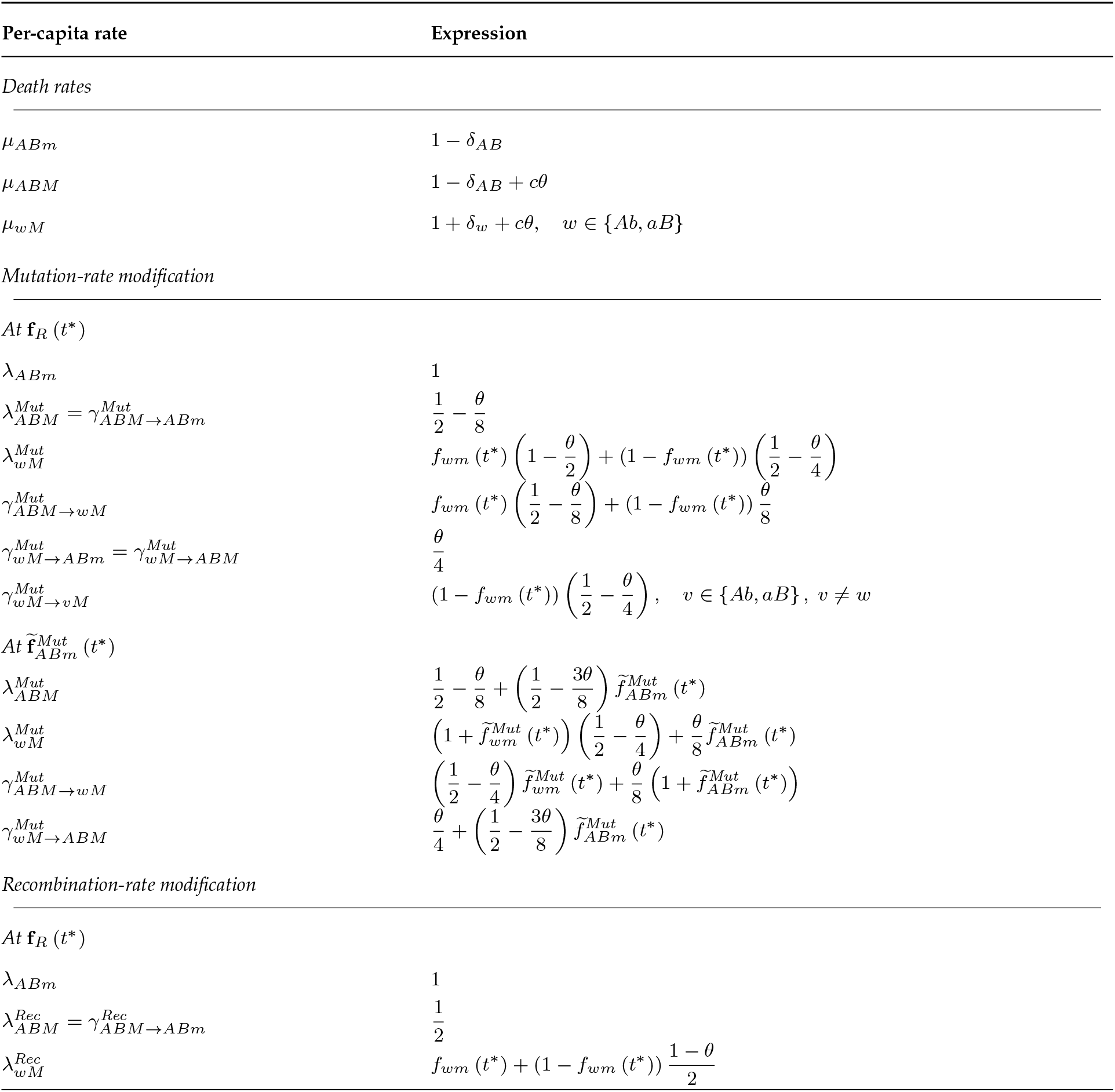
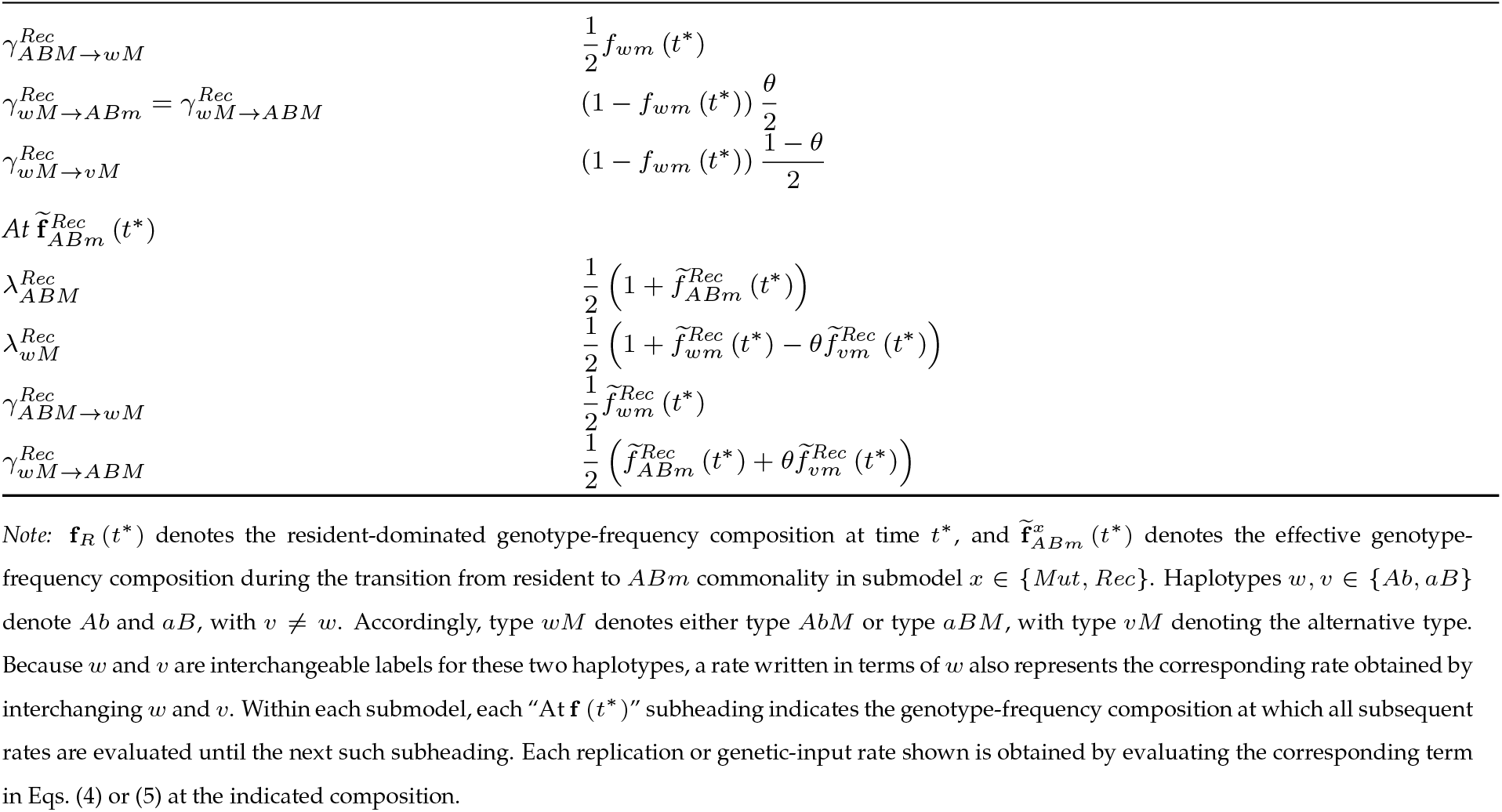
Per-capita rates used in the analytical approximations.

| Per-capita rate | Expression |
| --- | --- |
| *Death rates* |  |
| $\mu_{ABm}$ | $1 - \delta_{AB}$ |
| $\mu_{ABM}$ | $1 - \delta_{AB} + c\theta$ |
| $\mu_{wM}$ | $1 + \delta_w + c\theta, \quad w \in \{Ab, aB\}$ |
| *Mutation-rate modification* |  |
| $At \mathbf{f}_R(t^*)$ | |
| $\lambda_{ABm}$ | 1 |
| $\lambda_{ABM}^{Mut} = \gamma_{ABM \rightarrow ABm}^{Mut}$ | $\frac{1}{2} - \frac{\theta}{8}$ |
| $\lambda_{wM}^{Mut}$ | $f_{wm}(t^*) \left(1 - \frac{\theta}{2}\right) + (1 - f_{wm}(t^*)) \left(\frac{1}{2} - \frac{\theta}{4}\right)$ |
| $\gamma_{ABM \rightarrow wM}^{Mut}$ | $f_{wm}(t^*) \left(\frac{1}{2} - \frac{\theta}{8}\right) + (1 - f_{wm}(t^*)) \frac{\theta}{8}$ |
| $\gamma_{wM \rightarrow ABm}^{Mut} = \gamma_{wM \rightarrow ABM}^{Mut}$ | $\frac{\theta}{4}$ |
| $\gamma_{wM \rightarrow vM}^{Mut}$ | $(1 - f_{wm}(t^*)) \left(\frac{1}{2} - \frac{\theta}{4}\right), \quad v \in \{Ab, aB\}, v \neq w$ |
| $At \tilde{\mathbf{f}}_{ABm}^{Mut}(t^*)$ | |
| $\lambda_{ABM}^{Mut}$ | $\frac{1}{2} - \frac{\theta}{8} + \left(\frac{1}{2} - \frac{3\theta}{8}\right) \tilde{f}_{ABm}^{Mut}(t^*)$ |
| $\lambda_{wM}^{Mut}$ | $\left(1 + \tilde{f}_{wm}^{Mut}(t^*)\right) \left(\frac{1}{2} - \frac{\theta}{4}\right) + \frac{\theta}{8} \tilde{f}_{ABm}^{Mut}(t^*)$ |
| $\gamma_{ABM \rightarrow wM}^{Mut}$ | $\left(\frac{1}{2} - \frac{\theta}{4}\right) \tilde{f}_{wm}^{Mut}(t^*) + \frac{\theta}{8} \left(1 + \tilde{f}_{ABm}^{Mut}(t^*)\right)$ |
| $\gamma_{wM \rightarrow ABM}^{Mut}$ | $\frac{\theta}{4} + \left(\frac{1}{2} - \frac{3\theta}{8}\right) \tilde{f}_{ABm}^{Mut}(t^*)$ |
| *Recombination-rate modification* |  |
| $At \mathbf{f}_R(t^*)$ | |
| $\lambda_{ABm}$ | 1 |
| $\lambda_{ABM}^{Rec} = \gamma_{ABM \rightarrow ABm}^{Rec}$ | $\frac{1}{2}$ |
| $\lambda_{wM}^{Rec}$ | $f_{wm}(t^*) + (1 - f_{wm}(t^*)) \frac{1 - \theta}{2}$ |
| $\gamma_{ABM \rightarrow wM}^{Rec}$ | $\frac{1}{2} f_{wm}(t^*)$ |
| $\gamma_{wM \rightarrow ABm}^{Rec} = \gamma_{wM \rightarrow ABM}^{Rec}$ | $(1 - f_{wm}(t^*)) \frac{\theta}{2}$ |
| $\gamma_{wM \rightarrow vM}^{Rec}$ | $(1 - f_{wm}(t^*)) \frac{1 - \theta}{2}$ |
| $At \tilde{\mathbf{f}}_{ABm}^{Rec}(t^*)$ | |
| $\lambda_{ABM}^{Rec}$ | $\frac{1}{2} \left( 1 + \tilde{f}_{ABm}^{Rec}(t^*) \right)$ |
| $\lambda_{wM}^{Rec}$ | $\frac{1}{2} \left( 1 + \tilde{f}_{wm}^{Rec}(t^*) - \theta \tilde{f}_{vm}^{Rec}(t^*) \right)$ |
| $\gamma_{ABM \rightarrow wM}^{Rec}$ | $\frac{1}{2} \tilde{f}_{wm}^{Rec}(t^*)$ |
| $\gamma_{wM \rightarrow ABM}^{Rec}$ | $\frac{1}{2} \left( \tilde{f}_{ABm}^{Rec}(t^*) + \theta \tilde{f}_{vm}^{Rec}(t^*) \right)$ |

Note: $\mathbf{f}_R(t^*)$ denotes the resident-dominated genotype-frequency composition at time $t^*$ , and $\tilde{\mathbf{f}}_{ABm}^x(t^*)$ denotes the effective genotype-frequency composition during the transition from resident to $ABm$ commonality in submodel $x \in \{Mut, Rec\}$ . Haplotypes $w, v \in \{Ab, aB\}$ denote $Ab$ and $aB$ , with $v \neq w$ . Accordingly, type $wM$ denotes either type $AbM$ or type $aBM$ , with type $vM$ denoting the alternative type. Because $w$ and $v$ are interchangeable labels for these two haplotypes, a rate written in terms of $w$ also represents the corresponding rate obtained by interchanging $w$ and $v$ . Within each submodel, each “At $\mathbf{f}(t^*)$ ” subheading indicates the genotype-frequency composition at which all subsequent rates are evaluated until the next such subheading. Each replication or genetic-input rate shown is obtained by evaluating the corresponding term in Eqs. (4) or (5) at the indicated composition.

## Appendix E The effective genotype-frequency composition during the transition from resident to *ABm* commonality

To approximate the genotype-frequency composition encountered by type *ABM* and its descendants while an establishing *ABm* lineage increases from rarity toward commonality, we represent the transition by a single effective composition, 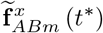. We first assign type *ABm* a pre-normalization weight, 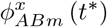, with the complementary weight 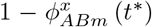 assigned to the two resident genotypes before accounting for their continued decline. Because the transition spans type-*ABm* frequencies from near zero to near one, we use its midpoint, 1*/*2, as a baseline for this weight and take its arithmetic mean with the probability that type-*wM* descendants of *ABM* regenerate haplotype *AB* after passing through intermediate genotypes. Under mutation-rate modification, either type *AbM* or type *aBM* is one mutation from *AB*, and regeneration requires mutation at the appropriate one of the two major loci. The regeneration probability associated with either path is therefore 1*/*2. Weighting these probabilities by the resident frequencies that determine the relative contributions of the two paths gives *f*_*Abm*_ (*t*^∗^) */*2 + (1 − *f*_*Abm*_ (*t*^∗^)) */*2 = 1*/*2, yielding

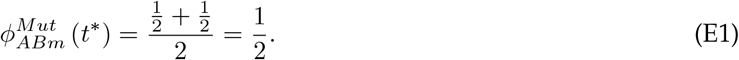

Under recombination-rate modification, type *AbM* regenerates *AB* by encountering type *aBm*, whereas type *aBM* does so by encountering type *Abm*. The corresponding regeneration probabilities are therefore 1 − *f*_*Abm*_ (*t*^∗^) and *f*_*Abm*_ (*t*^∗^). Weighting these probabilities by the resident frequencies that determine the relative contributions of the two paths gives the overall regeneration probability 2*f*_*Abm*_ (*t*^∗^) (1 − *f*_*Abm*_ (*t*^∗^)). Taking the arithmetic mean of this probability and the midpoint type-*ABm* frequency gives

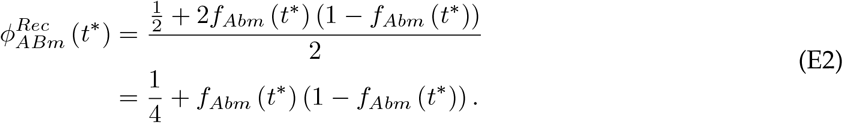

Thus, 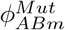 is independent of resident genotype composition, whereas 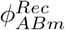 is largest when the complementary resident genotypes are equally frequent and decreases as their frequencies become asymmetric. The complementary weight, 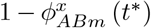, is assigned to the two resident genotypes according to their frequencies at time *t*^∗^. To account for their continued decline, each resident contribution is further multiplied by 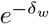, corresponding to its expected decline over one unit of model time. The resulting unnormalized contributions are therefore 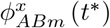 for type *ABm* and 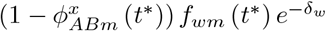 for type *wm*, whose sum gives the normalization factor

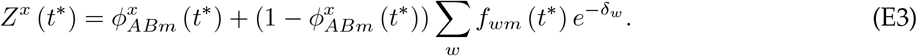

Normalizing the type-*ABm* contribution by *Z*^*x*^ (*t*^∗^) gives its effective frequency during the transition,

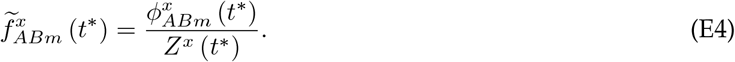

Likewise, normalizing each resident contribution by *Z*^*x*^ (*t*^∗^) gives its corresponding effective frequency,

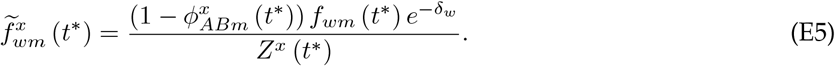

These normalized frequencies define the effective genotype-frequency composition encountered during the transition,

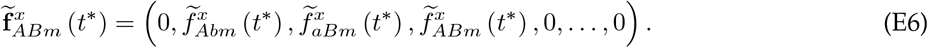

## Appendix F The appearance-weighted mean per-lineage establishment probability under stochastic resident genotype-frequency variation

Using diffusion theory (Waxman, 2011), we derive the appearance-weighted mean per-lineage establishment probability, 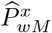, under stochastic resident genotype-frequency variation for the special case of equal resident mortality disadvantages, *δ*_*Ab*_ = *δ*_*aB*_ = *δ*_*R*_. We let *F* (*t*) = *F* denote the stochastic frequency of type *Abm*, with *dF* denoting its change over the infinitesimal time interval (*t, t* + *dt*). Approximating resident abundance at time *t* by its expectation, 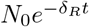, and noting that each resident organism replicates at percapita rate 1 and dies at per-capita rate 1 + *δ*_*R*_, the conditional first and second infinitesimal moments of *dF* under the diffusion approximation are

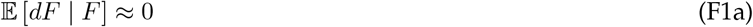

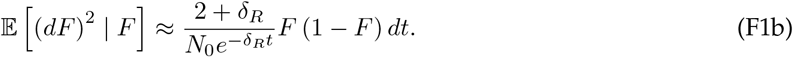

where E [· | *F* ] denotes expectation conditional on the current value of *F* (*t*). We define the rescaled time

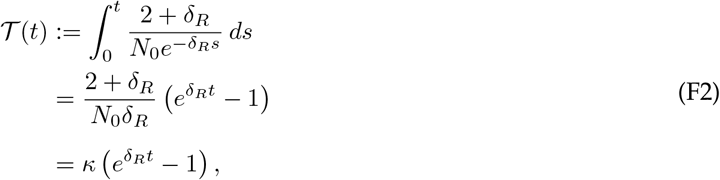

where *κ* := (2 + *δ*_*R*_)*/*(*N*_0_*δ*_*R*_) is the rescaling constant. Under this rescaling, the first three moment equations are *d*E [*F* ] */d*T = 0, *d*E *F* ^2^ */d*T = E [*F* ] − E *F* ^2^, and *d*E *F* ^3^ */d*T = 3 E *F* ^2^ − E *F* ^3^ . Because the frequencies obtained from expected resident abundances remain constant when *δ*_*Ab*_ = *δ*_*aB*_ = *δ*_*R*_, *F* (0) = *f*_*Abm*_ (0). Consequently, E [*F* ] = *f*_*Abm*_ (0), while

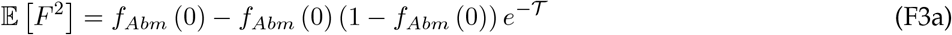

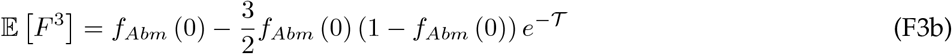

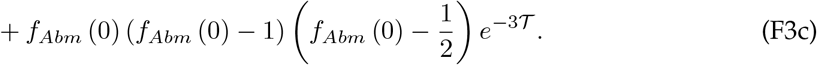

A type-*wM* lineage can arise through modifier-locus mutation only in a type-*wm* organism, so states with greater type-*wm* abundance contribute proportionally more type-*wM* lineage appearances. For a function *G*(*F* ), we denote expectation under the resulting appearance-weighted distribution by E_*w*_ [*G*(*F* )], where the subscript *w* identifies the type-*wM* lineage whose appearances determine the weighting. Since the frequencies of types *Abm* and *aBm* are *F* and 1 − *F*, respectively, and total resident abundance declines proportionally to 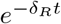, these expectations are

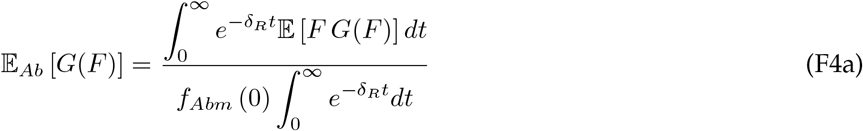

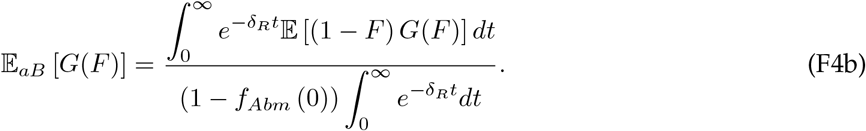

To average the exponential terms in Eq. (F3) over resident decline, we use the following identity for *ζ >* 0:

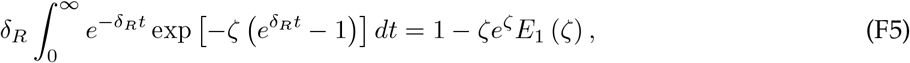

where the exponential integral, 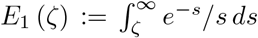, satisfies *E*_1_ (*ζ*) ≈ −Eu − ln *ζ* + *ζ* − *ζ*^2^*/*4 + *O ζ*^3^ for small *ζ*, with Eu ≈ 0.577216 denoting Euler’s constant. Retaining terms through third order gives 1 − *ζe*^*ζ*^*E*_1_ (*ζ*) ≈ X (*ζ*), where

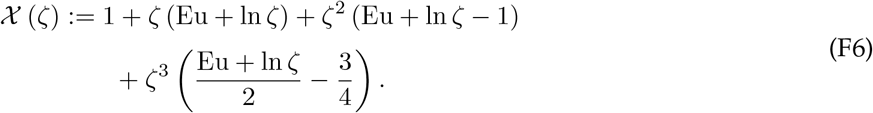

Since 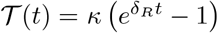, averaging the *e*^−*T*^ and *e*^−3*T*^ terms in Eq. (F3) gives X (*κ*) and X (3*κ*), respectively. Setting *G*(*F* ) = *F* in Eq. (F4) and substituting the moments in Eq. (F3), the appearance-weighted mean frequency of type *Abm* for type-*wM* lineage appearances is

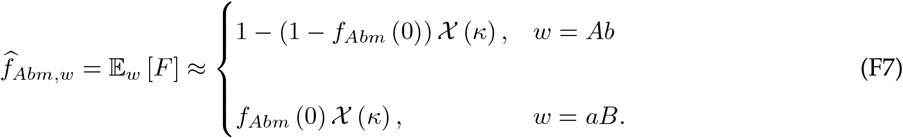

The corresponding appearance-weighted mean resident genotype-frequency composition is

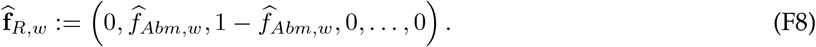

Similarly, setting *G*(*F* ) = *F* ^2^ in Eq. (F4), the corresponding appearance-weighted second moment is

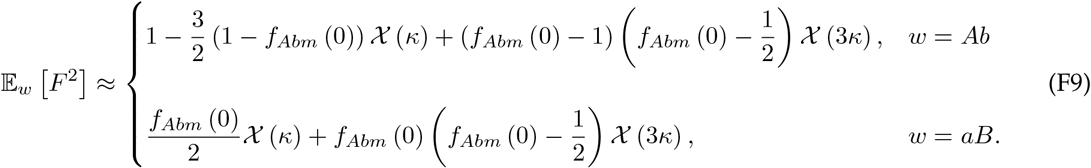

The corresponding appearance-weighted variance in type-*Abm* frequency, V_*w*_ := E_*w*_ *F* ^2^ − [E_*w*_ [*F* ]]^2^, is therefore

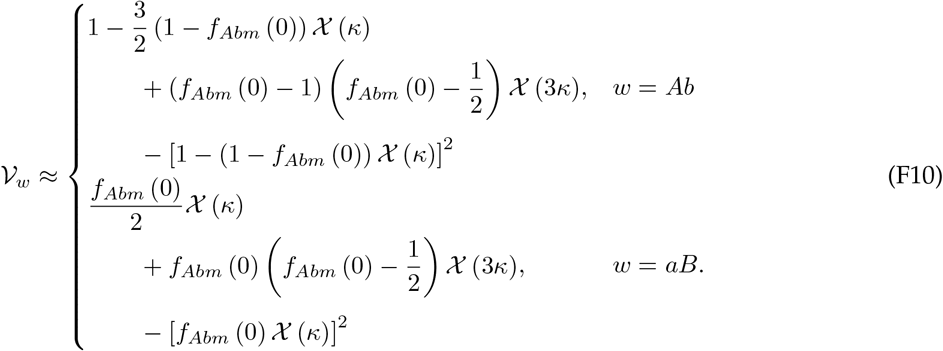

Because the resident genotype-frequency composition is completely determined by the frequency of type *Abm*, we define the establishment probability along this one-dimensional composition as 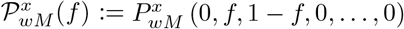. The desired appearance-weighted mean is then 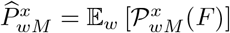. Expanding 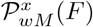 to second order around the appearance-weighted mean frequency, 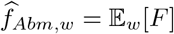, gives

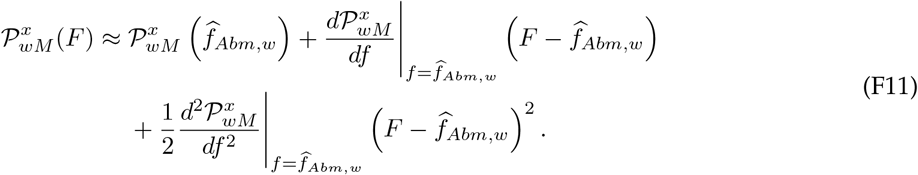

Taking the appearance-weighted expectation eliminates the first-order term because 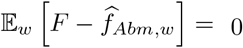, while 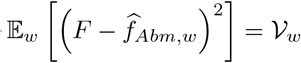. Neglecting third- and higher-order terms therefore gives the appearanceweighted mean per-lineage establishment probability adjusted for stochastic resident genotype-frequency variation:

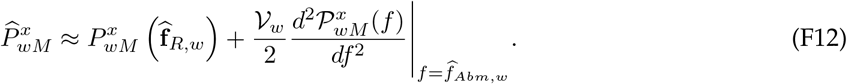

## Notes

### Competing Interest Statement

The authors have declared no competing interest.

